# A Generalized Platform for Artificial Intelligence-powered Autonomous Protein Engineering

**DOI:** 10.1101/2025.02.12.637932

**Authors:** Nilmani Singh, Stephan Lane, Tianhao Yu, Jingxia Lu, Adrianna Ramos, Haiyang Cui, Huimin Zhao

**Affiliations:** Carl R. Woese Institute for Genomic Biology, University of Illinois Urbana-Champaign, Urbana, IL 61801; Department of Chemical and Biomolecular Engineering, University of Illinois Urbana-Champaign, Urbana, IL 61801; NSF Molecule Maker Lab Institute, University of Illinois Urbana-Champaign, Urbana, IL 61801; DOE Center for Advanced Bioenergy and Bioproducts Innovation, University of Illinois Urbana-Champaign, Urbana, IL 61801

## Abstract

Proteins are the molecular machines of life with numerous applications in energy, health, and sustainability. However, engineering proteins with desired functions for practical applications remains slow, expensive, and specialist-dependent^1–3^. Here we report a generally applicable platform for autonomous protein engineering that integrates machine learning and large language models with biofoundry automation to eliminate the need for human intervention, judgement, and domain expertise. Requiring only an input protein sequence and a quantifiable way to measure fitness, this autonomous platform can be applied to engineer virtually any protein. As a proof of concept, we engineered *Arabidopsis thaliana* halide methyltransferase (*At*HMT)^4^ for a 90-fold improvement in substrate preference and 16-fold improvement in ethyltransferase activity, along with developing a *Yersinia mollaretii* phytase (*Ym*Phytase)^5,6^ variant with 26-fold improvement in activity at neutral pH. This was accomplished in four rounds over four weeks, while requiring construction and characterization of fewer than a total of 500 variants for each enzyme. This platform for autonomous experimentation paves the way for rapid advancements across diverse industries, from medicine and biotechnology to renewable energy and sustainable chemistry.

---

Laboratory experiments are essential to scientific research, which is traditionally driven by skilled researchers. However, many factors can slow down the conventional discovery process and limit efficiency. Manual laboratory tasks are time- and labor-intensive, prone to reproducibility issues, limited in scalability, and grow increasingly complicated when managing large datasets and high-throughput experiments. Autonomous experimentation has the potential to transform scientific research by enabling integration of artificial intelligence (AI) and robotics to iteratively propose hypotheses, design and conduct experiments, and refine models with minimal human intervention. AI-enabled systems can explore vast, multi-dimensional spaces more efficiently than traditional computational techniques while robotics and automation can perform experiments faster, more reliably and at higher throughputs with better scalability. Autonomous laboratories hold tremendous potential for diverse fields such as synthetic biology, chemical synthesis, and materials discovery by enabling more efficient and scalable research^7–9^. Early demonstration of such a system includes the Robot Scientist “Adam” that autonomously generated and tested functional genomics hypotheses for *Saccharomyces cerevisiae*^10^. Coscientist, a multi-LLMs-based intelligent agent, can autonomously perform chemical synthesis reactions^11^. In materials science, autonomous experimentation platform Ada optimized thin-film compositions^12^. The A-Lab combines robotics, machine learning (ML), and historical data to synthesize inorganic powders and successfully created 41 new compounds in 17 days of continuous operation^13^. A recent work developed autonomous mobile robots that can perform, analyze and decide next steps for exploratory synthetic chemistry reactions^9,14^.

In biology, autonomous experimentation is comparatively less mature, making even routine processes like DNA assembly, gene editing, or metabolic engineering a challenging task^7,15^. Moreover, integrating instruments for continuous experimentation requires skilled personnel with expertise at the intersection of biological experimentation, robotics, and programming^16–18^. Much previous work in autonomous synthetic biology has been highly specific, targeting singular goals like engineering a single protein^17^, metabolic engineering for a single product^19^, or orphan enzyme identification^10^. Despite these advances, a broadly applicable autonomous system must be highly generalizable for extensive utility. Generalizable platforms are more scalable and adaptable, able to address diverse problems across different locations without the need for new workflows. Widely adopted systems enable researchers to use a common scientific framework, promoting collaboration and efficient knowledge transfer while minimizing the need to redevelop similar methods for common goals. With scalable, generalizable platforms, synthetic biology can move beyond isolated successes and drive innovation in a wide range of fields.

In this work, we use protein engineering as a case study to establish a roadmap for generalized autonomous experimentation in synthetic biology. Protein engineering provides an extensive toolkit for modifying enzymes for widespread application in fields such as medicine, biofuels, and biocatalysis^1,2^. By iterative design, build, test, and learn (DBTL) cycles, enzymes can be made more stable, selective, or efficient. Methods like directed evolution and computer-aided design offer diverse strategies for protein engineering and high-throughput screening strategies allow iterating over a large sample space^3,20,21^. Despite the wide applicability of protein engineering, there remain unmet needs, particularly in efficiently navigating vast sequence spaces and optimizing protein function in complex environments. Autonomous protein engineering represents the state-of-the-art in addressing these challenges through self-driving frameworks for executing the DBTL cycle^16,22,23^. Many synthetic biology applications have demonstrated decent success for autonomous design, such as in metabolic engineering^19^ and construction of plasmids^24^. A recent study^17^ demonstrated an automated protein engineering campaign that improved glycoside hydrolase thermostability using Bayesian optimization, cell-free expression and gene fragment-based variant creation^17^. However, reliance on an external cloud lab adds a layer of opaqueness while the high cost of gene fragments, limitations of cell free systems, and enzyme-specific generation of variant libraries reduce the workflow’s overall versatility and scalability.

Here, we present a generalized platform for autonomous protein engineering enabled by the Illinois Biological Foundry for Advanced Biomanufacturing (iBioFAB) (Extended Data Fig. 1), ML, and large language models (LLMs) (Fig. 1, Extended Data Fig. 2). The process begins by designing a high quality mutant library using a protein LLM^25^ and epistasis model^26^. The library is constructed and screened by iBioFAB using optimized modular and integrated workflows. The assay data from each cycle is used for training a low-N ML model^27^ to predict variant fitness for subsequent iterations. Since this platform requires only a protein sequence and quantified fitness data, it can be applied to virtually any protein. As a proof of concept, in four rounds within four weeks, we engineered variants of two enzymes, *Arabidopsis thaliana* halide methyltransferase (*At*HMT)^28^ and *Yersinia mollaretii* phytase (*Ym*Phytase)^5^ with approximately 16- and 26-fold higher activity compared to the wild type enzymes, respectively.

**Figure 1:**
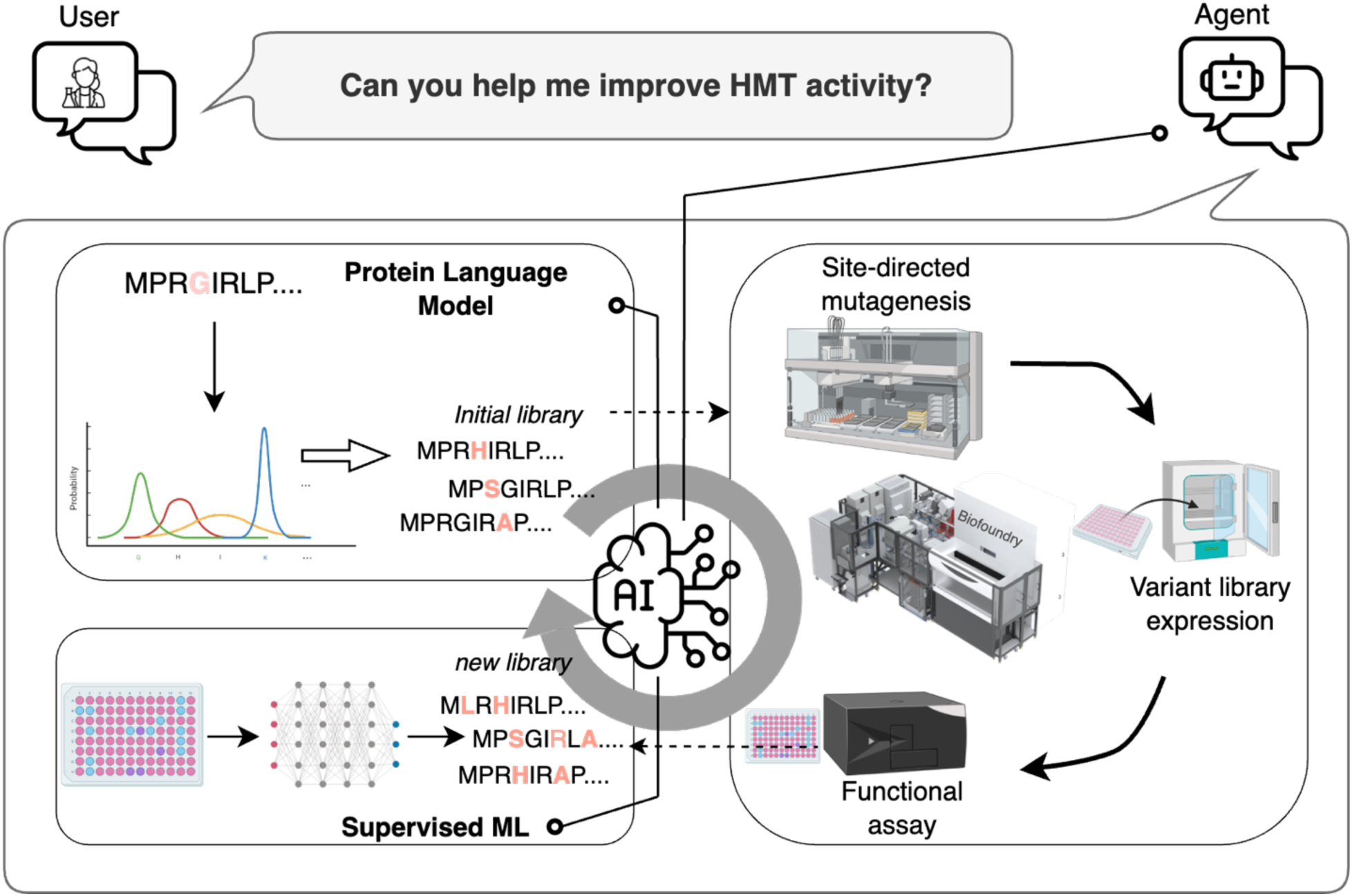
Overview of the generalized platform for autonomous protein engineering. The platform consists of three modules. First, a sequence-based unsupervised predictive model generates variants for the initial library. This variant library is created by an integrated biofoundry, a laboratory automation platform that combines various instruments for robust and continuous experimentation. Next, the biofoundry measures the fitness of the variants, and the resulting data is used to train a supervised ML model to predict subsequent variants. The seamless integration of protein language models, automated experimentation, and ML minimizes the need for human decisions and interventions during the protein engineering process. This autonomous protein engineering platform can design, test, and create improved enzymes without input from a human operator.

## Automating construction and characterization of protein variants on a biofoundry

The iBioFAB (Extended Data Fig. 1) has been used to automate various biological processes, such as pathway optimization^19^, yeast genome engineering^29^, TALEN assembly^30^, plasmid design and construction^24^, and natural product discovery^31,32^. In ML-guided protein engineering, sequencing and verifying the mutants generated through site-directed mutagenesis (SDM) often delays the process and increases costs. To design a robust and continuous workflow, we developed a HiFi-assembly based mutagenesis method (Fig. 2B, Extended Data Fig. 3) that eliminated the need for sequence verification in the middle of the protein engineering campaign, enabling an uninterrupted workflow. Throughout multiple rounds of protein engineering, some mutants were randomly selected, sequenced, and confirmed to have the correct targeted mutations with around 95% accuracy (Fig. 2C). All higher-order mutants are combinations of single mutants from the initial library, with one additional mutation added in each round. Variants with multiple mutations can be generated through SDM of a template plasmid containing one fewer mutation and, as such, there is no need to order new primers for each iterative cycle, thereby saving time and cost. This optimized high-fidelity approach was crucial for a reliable and continuous workflow during iterative cycles of protein engineering.

**Figure 2:**
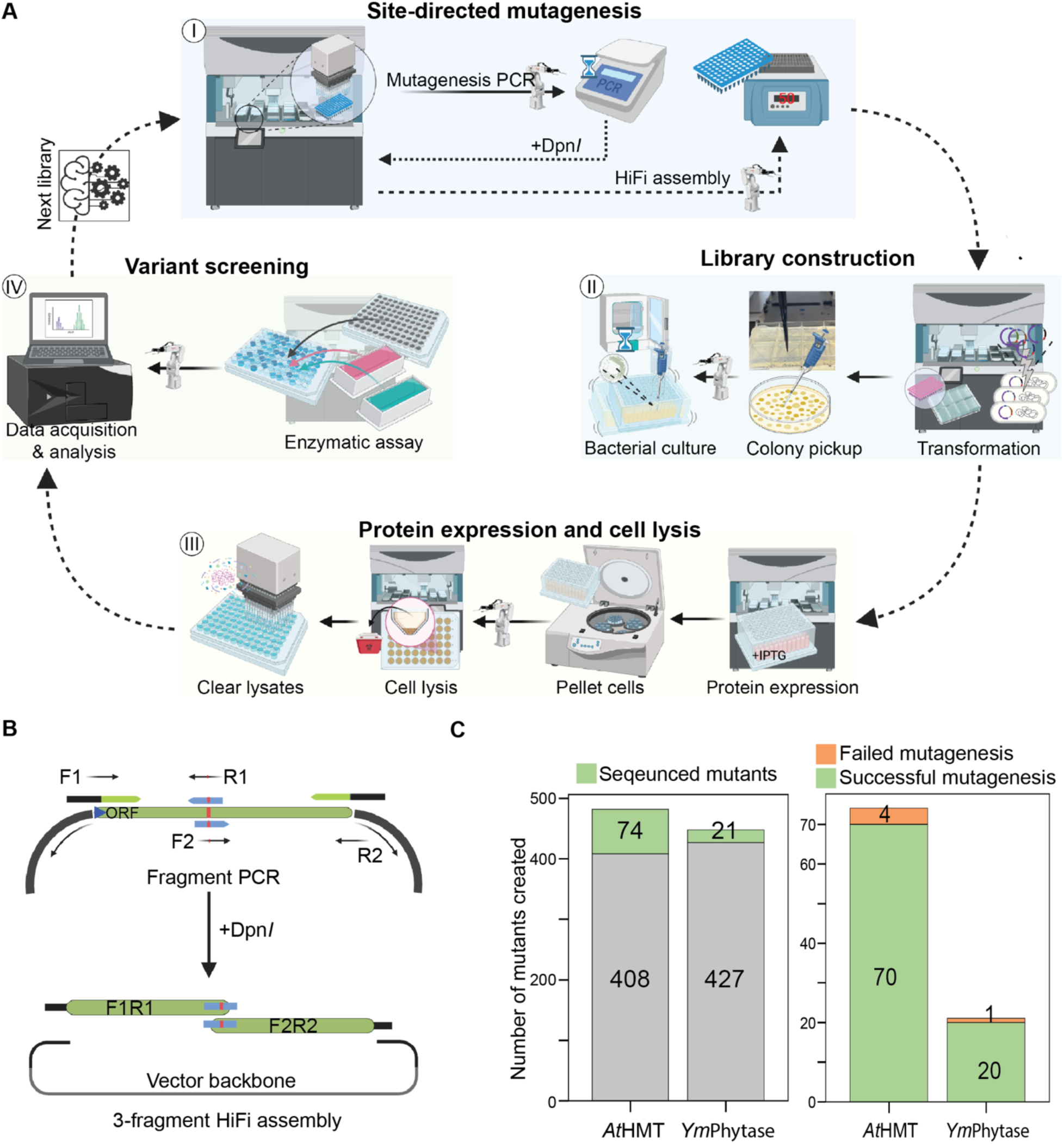
Automated Protein Engineering Workflow. **(A)** Overview of the laboratory automation pipeline used for autonomous protein engineering. The workflow was divided into smaller modules, programmed into iBioFAB instruments, and optimized for robustness. The continuous workflow for a complete round of screening, from mutagenesis PCR to functional assay, is completed within seven days (Extended Data Figs. 4-8). **(B)** A PCR and HiFi based site-directed mutagenesis protocol was optimized for fidelity. The mutagenesis PCR generates two DNA fragments, which are then assembled with the vector backbone using HiFi assembly. **(C)** We created 482 mutants for *At*HMT, and DNA sequencing confirmed that 70 of 74 variants had the correct mutations. For *Ym*Phytase, we created 448 mutants and 20 of 21 were correctly mutated based on sequencing. Overall, the success rate for mutagenesis was about 95% when picking only a single colony for each variant.

The protein engineering workflow was divided into seven automated modules for robustness and ease of troubleshooting (Fig. 2A, Extended Data Fig. 4), allowing recovery without restarting the entire process. Each instrument on the iBioFAB was programmed for specific tasks, scheduled via Thermo Fisher *Momentum* software, and fully integrated by a central robotic arm. The robotic pipeline automates all protein engineering steps including mutagenesis PCR, DNA assembly, transformation, colony picking, plasmid purification, protein expression, and enzyme assays (Extended Data Figs. 5-8).

## Design of protein variants using protein LLM and ML

The design of the initial library is crucial to the protein engineering campaign as a diverse and high-quality library increases the likelihood of efficient optimization^33^. To create a generally applicable autonomous protein engineering platform, we employed a combination of a protein LLM and an epistasis model. This approach maximized both the diversity and quality of the initial library, enhancing the chances of identifying promising mutants early in the process. We used a state-of-the-art protein LLM, ESM-2^25^, a transformer model trained on global protein sequences. ESM-2 predicts the likelihood of amino acids occurring at specific positions based on sequence context, and the likelihood can be interpreted as variant fitness^34^. Additionally, we included an epistasis model, EVmutation^26^ focusing on the local homologs of the target protein.

To demonstrate the generality of our autonomous protein engineering platform, we selected two distinct enzymes, *At*HMT and *Ym*Phytase, with a goal of improving two different enzymatic properties. The halide methyltransferase *At*HMT offers potential for synthesizing *S*-adenosyl-L-methionine (SAM) analogs, which are essential for biocatalytic alkylation, a process with crucial biological applications^28,35^. However, the chemical synthesis and commercial availability of SAM analogs are often limited. *At*HMT has shown promising promiscuous alkyltransferase activity, allowing for the synthesis of SAM analogs from more readily available and cost-effective alkyl halides and *S*-adenosyl-L-homocysteine (SAH)^4,36^. We focused here on improving the ethyltransferase activity of *At*HMT, seeking to improve its preference for ethyl iodide over methyl iodide. We secondly aimed to broaden the pH range for activity of *Ym*Phytase, a phosphate-hydrolyzing enzyme that exhibits low activity at neutral pH^37^. Phytases can hydrolyze phytic acid, the main storage form of phosphate in plant tissues that is indigestible by most animals, and release inorganic phosphate, an important nutrient^38^. However, to function in animal feed, phytase must have high activity at a broad pH range to tolerate the pH variation of the gastrointestinal tract of animals. Most phytases function optimally in acidic conditions and exhibit a sharp loss of activity at neutral pH, motivating the development of phytase enzymes with improved activity closer to neutral pH. We therefore engineered *Ym*Phytase, which has better activity than commonly used fungal phytases and has shown promising improvements in previously reported directed evolution campaigns^5,6^.

By combining the two unsupervised models, we generated a list of 180 variants each of *At*HMT and *Ym*Phytase for initial screening (Fig. 3A-B, Extended Data Fig. 2). We found that 59.6% of *At*HMT and 55% of YmPhytase variants performed above the wild type baseline (Fig. 3C, Extended Data Figs. 9, 12), with 50% and 23% being significantly better, respectively (two-tailed Student’s t-test *p* < 0.05, Extended Data Figs. 9, 12). The mutants generated by EVmutation performed better than those predicted by ESM-2, although there was significant overlap between predictions of the two models (Extended Data Fig. 29). The best mutants from the unsupervised models’ predictions were V140T for *At*HMT, with a 2.1-fold improvement, and V141M for *Ym*Phytase, with a 2.6-fold improvement. For *At*HMT, both models had independently identified V140T as one of the top ranked variants, which was reported as the best mutant in a previous semi-rational design study^4^. Whereas, for *Ym*Phytase, the predictions from both models did not include previously identified sites T44 and K45 in top predictions ^5^.

**Figure 3:**
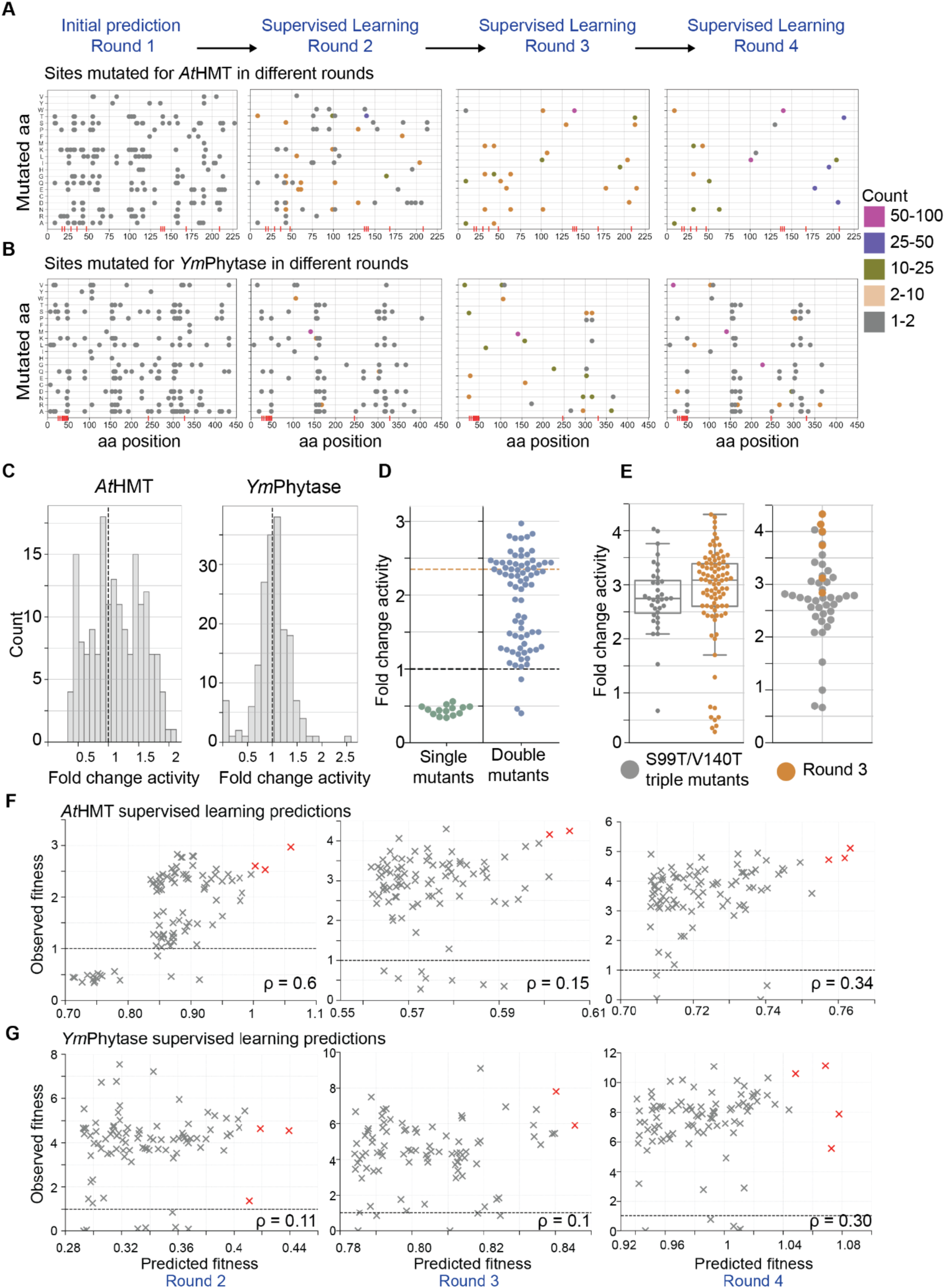
ESM-2, EVmutation, and low-N supervised learning ML models predict better variants during the screening of *At*HMT and *Ym*Phytase. **(A-B)** The individual mutations predicted in each round at various sites of the protein are marked and color coded by their prevalence in the screened variants. The initial sequence-based models start with a wide search, followed by narrowing down by a supervised learning model to a few select mutation sites in later rounds. The active site for *At*HMT (P20, V23, L27, W36, W47, Y139, V140, C143, Y172, R214) and *Ym*Phytase (R37, R41-T47, E241, D327) are marked by red lines on x-axis. The x-axis represents the mutation positions, and the y-axis represents the mutated amino acids. The exact amino acid sequence for both is available in the Supplementary Materials. **(C)** The majority of variants predicted by unsupervised models for the first round show higher activity than the wild type for both *At*HMT and *Ym*Phytase. The black dashed line shows wild-type activity. **(D)** None of the new single mutants predicted for the second round of *At*HMT engineering show better activity than wtHMT. Black dashed line indicates WT activity and orange dashed line shows V140T activity. **(E)** None of the S99T/V140T-containing triple mutants show better activity than the top predicted triple mutants from the third round. A comparison between 36 S99T/V140T-containing triple mutants with all the third-round mutants (right). Comparison of activities for six of the best predicted third round mutants were run in the same assay plate as the 36 triple mutants containing S99T/V140T mutation (left). **(F-G)** The predicted fitness by the low-N supervised learning model and observed fitness of mutants for *At*HMT and *Ym*Phytase. The dashed black line indicates the wild-type activity for respective proteins. Red crosses indicate mutants with the best predicted fitness according to the supervised model. The Spearman rank correlation coefficient is displayed at the bottom of the graphs.

Starting from the second round, the data from all previous rounds were used to train a supervised low-N regression model^27^. Once trained, the model predicted variants with one additional mutation than the previous round for the subsequent screening cycle (Extended Data Fig. 2). While the unsupervised models started with a wide search space in the first round, the supervised learning model narrowed to focus on a few key mutations in the top 90 predictions for both proteins (Fig. 3A-B). We constructed the top 96 predicted mutants and 90 were used for enzyme activity screening along with controls. Although there was little overall consistency between observed enzyme activities and the predictions of the low-N model, the model was nonetheless able to predict a subset of mutants with substantially improved activity in each round of engineering (Fig. 3F-G). For the second round of *At*HMT engineering, we predicted additional single mutations after training the low-N model using the data from the first round. The 14 new single mutant predictions performed poorly compared to the wtHMT (Fig. 3D, Extended Data Fig. 9) suggesting that the low-N model may not be well suited for finding new single mutation variants as compared to the unsupervised models used in the first round.

## New insights through ML-guided protein engineering

ML-driven experimentation often leads to surprising results and unexpected directions, as shown by the model’s prediction of counter-intuitive variants during the third round of *At*HMT engineering. None of the S99T/V140T-containing triple mutants were ranked in the top 100, despite S99T/V140T being the best double mutant from the second round. We tested the top 36 ranked triple mutants containing S99T/V140T and compared them against the model’s top predictions for triple mutants. Most S99T/V140T-based triple mutants failed to significantly improve activity over the V140T mutant with only 4/36 (11%) showing significantly better activity than V140T (two-tailed Student’s t-test *p* < 0.05, Fig. 3E, Extended Data Fig. 10), whereas the model-predicted triple mutants showed superior performance with 74/90 (82%) performing significantly better than V140T (two-tailed Student’s t-test *p* < 0.05, Extended Data Fig. 19). This suggests the ML model can potentially recognize synergistic effects between mutations, predicting more effective combinations than human-designed mutants.

We observed an increase in the desired activity throughout all four engineering cycles (Fig. 4A-D, Extended Data Figs. 9, 12). The fitness of variants improved in each round compared to the wild type for both *At*HMT and *Ym*Phytase (Fig. 4A-D). In four rounds of protein engineering, we screened 446 variants of *At*HMT and 448 variants of *Ym*Phytase using a crude lysate assay. For *At*HMT, the top performing variant in each subsequent round was 2.1-, 3-, 4.3- and 5.1-fold better than wtHMT in the crude lysate assay (Extended Data Figs. 9, 11). For *Ym*Phytase, the top performing variant in each subsequent round was 2.6-, 7.5-, 9- and 11.1-fold better than wtPhy in the crude lysate assay (Extended Data Fig. 12). We purified some of the best performing mutants from each round and characterized the enzyme kinetics (Extended Data Fig. 17-18). The best identified *At*HMT variant (round 4: V140T/L101I/E206D/S63N) showed an approximately 16-fold improvement in ethyltransferase activity compared to wtHMT (Fig. 4E, Extended Data Fig. 15-16, Extended Data Table 3). Another *At*HMT variant (round 4: V140T/L101I/T213S/V178E) showed an approximately 90-fold increase in preference for ethyl iodide over methyl iodide (Extended Data Table 1-2, 4).

**Figure 4:**
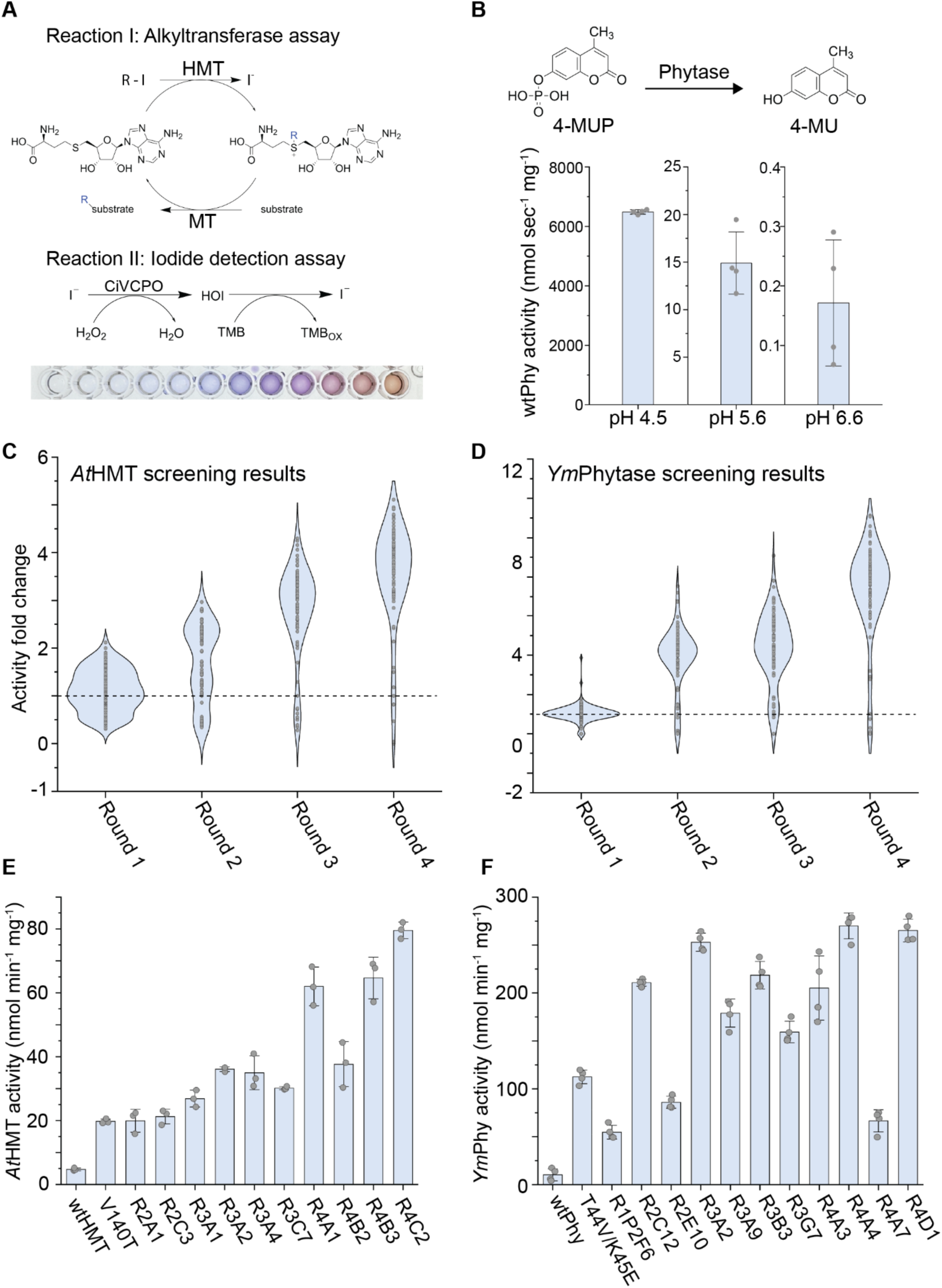
Results from AI-powered autonomous engineering of *At*HMT and *Ym*Phytase proteins. **(A)** Two sequential reactions were used to quantify the activity of *At*HMT: (I) the HMT-catalyzed release of iodide from ethyl iodide and SAH and (II) colorimetric quantification of iodide by measuring absorbance at 570 nm. **(B)** The phytase activity assay measured 4-MUP hydrolysis to 4-MU by monitoring fluorescence (*λ*_ex_ = 354 nm / *λ*_em_ = 465 nm). The wild type exhibits a sharp decrease in activity at pH 6.6. **(C)** The fold change in HMT ethyltransferase activity with respect to the wild type is plotted as violin plot for four rounds of crude lysate screening. **(D)** The fold change in phytase activity with respect to the wild type is plotted as violin plot for four rounds of crude lysate screening. (**E-F**) A selection of the best performing mutants from each screening round were purified and specific activity was calculated at 15 mM ethyl iodide for *At*HMT (n = 3) **(E)** and at 0.9 mM 4-MUP for *Ym*Phytase (n = 4) **(F)**. R1P1 and R1P2 denote plates 1 and 2 respectively from the first cycle of protein engineering. Labels with R2, R3, and R4 denote subsequent screening rounds (e.g. R2A1 denotes well A1 from the second round). The enzyme activity assay during screening used n ≥ 3 replicates. The mutations present in each variant can be found in Extended Data Tables 3 and 5. Full kinetic characterization of the *At*HMT (Extended Data Fig. 14-16) and *Ym*Phytase (Extended Data Figs. 22-28) mutants are provided as extended data figures.

For *Ym*Phytase, one variant from the fourth round (round 4: V141M/K226G/I15V/Q362R) exhibited a 26.3-fold improvement in specific activity over wtPhy at pH 6.6 (Fig. 4F). Another variant (round 4: V141M/K226G/I15V/E316P) performed remarkably well at both pH 5.6 and 6.6, exhibiting 25.8- and 19.8-fold improvements in specific activity compared to wtPhy, respectively (Fig. 4F, Extended Data Fig. 20). All proteins selected for purification showed a higher specific activity (Extended Data Table 5) and improved kinetic parameters (Extended Data Table 6) compared to the wild type at pH 4.5, 5.6, and 6.6. Since the screening involved several hours to obtain complete data from multiple replicates, the cell lysates contained a protease inhibitor cocktail in the crude lysates. After purification of selected phytase variants we noticed that the addition of the protease inhibitor cocktail, and more specifically one of its components 4-(2-aminoethyl) benzenesulfonyl fluoride hydrochloride (AEBSF), was critical to observe high activities of *Ym*Phytase enzyme variants (Extended Data Fig. 21). Screening conditions are therefore crucial to the design of the protein engineering campaign as proteins are optimized for the specific assay environment.

In the previous work that identified V140T as an improved variant^4^, 10 residues in the active sites were screened using NNK site-saturation libraries. Further screening by generating double and triple mutant libraries combining active site residues didn’t yield a better mutant than V140T^4^. In the current work, V140T is present in all third and fourth round variants and is the only mutation in the active site. All other mutation sites in top performing mutants (L101, T213, V195, E206, D9, E32) from the fourth round of engineering are located relatively far from the active site and on the surface of the protein (Movie S1). Moreover, the majority of fourth round variants show significantly improved activity over V140T alone.

In contrast, our *Ym*Phytase engineering yielded a new mutation V141M during the first round of screening that significantly improved activity compared to the wild type and was carried over into all subsequent iterative cycles. None of the top predictions by the unsupervised models included the previously identified T44/K45 sites discovered by semi-rational design^37^. Since the *Ym*Phytase crystal structure was not available, it was predicted using AlphaFold^39^. The majority of variants in the fourth round contain three mutations V141M, K226G, and I15V, of which V141 and K226 residues are close to the active site, while I15 is located at a more distant flexible region. Most other common mutation sites in the fourth round such as Q167, T25, L303 are distant from the active site, while A65 is located close to the active site (Movie S2).

## Language model-based user interface

As mentioned above, we demonstrated that using unsupervised protein mutation prediction tools and supervised regression models can significantly speed up protein engineering without requiring domain expertise in rational design. To further simplify access, we developed a natural language-based user interface utilizing LLMs (28) (Fig. 1). This allows users without programming skills to easily interact with our protein engineering platform. We created a user interface, leveraging OpenAI’s assistant API with customized functions, to design the initial library using simple language commands, like “help me improve phytase” or “design the initial library for *At*HMT”, especially for users without required coding skills. The system responds by accessing pre-programmed functions to execute sequence-based predictions and train regression models, making the platform accessible to non-technical users.

## Discussion

Through systematic development and optimization, we have demonstrated a generally applicable autonomous protein engineering platform which accepts a protein sequence as the sole input and requires minimal human decision-making or specialized protein knowledge. Without the need for human judgement, the autonomous platform is fully scalable and compatible with virtually any proteins having a quantitative protein function assay. Our natural language user interface further reduces barriers to entry, allowing users to operate the platform without specialized coding knowledge. We used two comprehensive case studies, *At*HMT and *Ym*Phytase, to demonstrate the autonomous protein engineering platform in action. Starting with only the wild type sequences, our approach autonomously performed successful engineering campaigns for both proteins by identifying variants with 16-fold improvements for *At*HMT and 26-fold improvements for *Ym*Phytase within four iterative cycles of protein engineering.

A few previous attempts have prototyped automated experimentation using Bayesian optimization for traversing a relatively limited search space^17,19,40^ for protein engineering. With the unimaginably vast and high-dimensional sample space of proteins, state-of-the-art AI tools are better suited for efficient exploration^41^. A recent study^17^ used a cell-free expression system, which is challenging to generalize due to many issues such as limited protein yield, poor protein folding, instability and degradation. That work required 20 iterative cycles to identify the final improved variants^17^, whereas our framework needed only four rounds to generate variants with substantial improvement. In many protein engineering campaigns, initial variant libraries are tailored for proteins of interest and require deep expertise about structure and function of the target. In contrast, we present a generalizable framework that can predict variants with only the protein amino acid sequence for the initial library and fitness data for iterative engineering cycles. This coupled with our stepwise approach for mutagenesis offers a broadly applicable and cost-effective solution for autonomous protein engineering.

The modularity of our framework makes possible substantial expansions and improvements in the future. With advancements in the state-of-the-art AI models for protein engineering, we expect this framework to improve as new models can be integrated into this platform. The exploration of higher-order mutants with substantially greater numbers of mutations becomes feasible with this platform, a challenge that has proven difficult to address especially for *in vitro* protein engineering studies. Moreover, while we used crude lysate assays in this study, this framework is compatible with many quantified measurements of protein activity, including *in vitro*, *in vivo*, growth-coupled, or reporter-based assays. By incorporating this flexibility, we have developed a general-purpose autonomous protein engineering platform that maximizes its potential applications.

Autonomous experimentation platforms have the potential to transform protein engineering and related fields including synthetic biology^42^, biocatalysis^43^, metabolic engineering^44^, and retrobiosynthesis^45^. The rapid progress of laboratory automation, such as liquid-handling robots^46^, microfluidics^47^, laboratory virtualization services^48^, and increasing global investment into biofoundries^49^ will make these platforms more accessible to researchers. The integration of ML tools and robotic laboratory platforms offers round-the-clock operation, improved reproducibility, and increased efficiency while minimizing the need for human decision-making. Various applications, including the design of new-to-nature enzymes^50^, discovery of novel natural products^31,32^, enhancement of drug and biomolecule efficacy, and the optimization of products for sustainable chemistry^43^, stand to benefit greatly from the transformative power of autonomous experimentation. The broadly applicable framework presented here for protein engineering forms a crucial component of the autonomous research toolbox, establishing a strong foundation for the future of synthetic biology.

## Methods

### LLM and supervised learning for prediction of protein variants

Two different zero-shot prediction models were used to guide the design of the initial variant library: EVmutation and ESM-2. EVmutation is a statistical model designed by Hopf et al.^26^ It captures the co-variations between pairs of residues in an amino acid sequence. This is achieved by fitting a pairwise graph model to the multiple sequence alignment (MSA) of the homologous to the target protein. The model then scores the impact of amino acid substitutions by calculating the log-ratio of sequence probabilities between the mutant and wild-type sequence. In this work, we utilized the integrated web server EVcouplings ^51^ with the default searching parameters and highlighted bit-scores. ESM-2 is a transformer-based protein language model designed by Rives et al.^52^ It is trained on large and diverse protein sequence database and captured the rules governing the protein structure and functions. ESM-2 can output the probability of a certain amino acid occurring at a given position based on the up and down stream context. Thus, we can score any given mutation by using wild-type as a reference and compare the probability of the given mutation with that of the wild-type. In this work, we used the workflow implement in ESM-2 (https://github.com/facebookresearch/esm) to make the prediction with the model esm2_t36_3B_UR50D.

For each round of engineering, we trained a supervised prediction model based on all experimentally measured variant fitness data from current round and all rounds prior. We first preprocess raw data by removing all zero or negative variants and normalize the data by taking the log scale. In this work, we trained the supervised model by modifying the workflow implement by Hsu et al.^27^ (https://github.com/chloechsu/combining-evolutionary-and-assay-labelled-data).We initially trained the model using augmented Potts and ESM approach, but the evolutionary density score feature has negatively correlated with experimental results. Thus, we modified the original training script and used the command ‘python src/evaluate.py phytase onehot --n_seeds=1 --n_threads=1 --n_train=-1’ for training and prediction.

### GPT-based user interface

To implement a Generative Pre-trained Transformer (GPT)-based user interface, OpenAI’s assistant Application Programming Interface (API) (https://platform.openai.com/docs/assistants/overview) was employed. The assistant was configured using the ‘gpt-4-turbo’ model. The user interface carries tools of file-search (https://platform.openai.com/docs/assistants/tools/file-search) and function-calling (https://platform.openai.com/docs/assistants/tools/function-calling). The user interface will invoke file searching to respond to user’s general questions and invoke function calling to assist the design of initial variant library.

### Expression plasmids

The HMT and CiVCPO plasmids were a gift from Uwe Bornscheuer lab. Both the HMT plasmids and empty vector (pET-28a) were transformed in *Escherichia coli* BL21 (Δmtn) strain, a gift from Uwe Bornscheuer lab, which is an MTA/SAH nucleosidase knockout *E. coli* BL21(DE3) strain^4^. The phytase cDNA was ordered from Twist Biosciences and cloned in pET-28a vector using HiFi assembly. These plasmids were sequence verified using Plasmidsaurus (San Francisco, CA, USA).

### Site-directed mutagenesis

For site-directed mutagenesis, three PCR fragments were assembled using HiFi assembly (Extended Data Fig. 1). The mutagenesis primers (27 base-pairs, 12+3+12) were designed using a Python script and ordered in 96-deepwell plates from IDT. For PCR templates, the plasmids were linearized with restriction enzymes and purified using a PCR cleanup kit (Zymo #D4018). The HMT plasmid was linearized with *EcoR*V (for mutagenesis PCR of the ORF) and *Xba*I (for backbone PCR). The phytase plasmid was linearized with *Hpa*I (for mutagenesis PCR of the ORF) and *Xho*I (for backbone PCR). The PCR fragments contained 27-40 overlapping base pairs. The PCR for the vector backbone was treated with *Dpn*I (37°C for four hours, followed by overnight incubation at room temperature) and purified using a PCR cleanup kit (Zymo #D4018), and stored at −20 °C until used. The mutagenesis PCR for forward and reverse fragments was run in separate 96-well plates, followed by *Dpn*I treatment. Q5 DNA polymerase (NEB #M0491) was used for mutagenesis PCR (50 µL reaction, ∼200 pg template, 18 cycles, 65 °C 30 seconds, and 72 °C one minute). After PCR, 2.5 µL of the reaction was mixed with 50 µL of 1x EvaGreen Dye (Biotinum #31000) and fluorescence (*λ*_ex_ = 488 nm / *λ*_em_ = 535 nm) was measured to verify successful PCR.

HiFi assembly reactions (15 µL) were prepared using 30 ng of purified vector backbone PCR, 1.25 µL each of *Dpn*I treated forward and reverse PCR reactions, and 7.5 µL of HiFi master mix (NEB #E2621) followed by incubation at 50 °C for 30 minutes. To increase the efficiency of HiFi assembly, 50ng of single-strand binding protein (NEB #M2401) was added to 500 µL HiFi master mix^53^. Homemade competent DH5α cells in a 96-well plate (Biorad # HSS9641) were transformed with 5 µL of the HiFi reaction by 30 seconds heat shock, followed by one hour outgrowth in 150 µL SOCS media. DH5α outgrowth cultures were spread on 8-well omnitray plates (120 µL per well, kanamycin 50 µg/mL) and incubated overnight at 37 °C. DH5α colonies were picked using Pickolo and incubated overnight at 37 °C in a 96-deepwell plate containing 1 mL of Terrific Broth (TB) and kanamycin. Minipreps were then performed using PureLink Pro Quick96 Plasmid Purification Kit (Invitrogen #K211004A). Miniprep DNA of some mutants from the 96-well plate was sent for sequencing. The miniprep DNA was used to transform competent BL21 cells in a 96-well plate for further protein expression and functional assays.

### *E. coli* heat shock competent cell preparation in 96-well plate

DH5α, BL21, and BL21(Δmtn) competent cells were prepared in the lab and stored in a high-profile 96-well PCR plate (Bio-Rad #HSS9641) at −80°C. Briefly, an overnight preculture (no antibiotics, 37 °C) was started from a single colony. A 250 mL LB media was inoculated with 5 mL of the starter culture and grown at 30 °C until the OD reached 0.4–0.6, and then transferred to cold room (4 °C) for one hour. The cells were centrifuged at 1000xg for 20 minutes at 4 °C. The cells were then gently resuspended in Buffer RF1 (100 mM rubidium chloride, 50 mM manganese chloride, 30 mM potassium acetate, 10 mM calcium chloride, 15% w/v glycerol, pH 5.8), incubated at 4 °C for 30 minutes, then centrifuged. Subsequently, the cells were resuspended in 10 mL of RF2 buffer (10 mM MOPS, 10 mM rubidium chloride, 75 mM calcium chloride, 15% w/v glycerol, pH 5.8). 50 µL was aliquoted into each well of a 96-well plate in a cold room, followed by sealing and snap-freezing in liquid nitrogen. The cells were stored at −80 °C until used.

### Iodide detection assay for the ethyltransferase activity of HMT

We followed the iodide quantification protocol developed by Tang et al.^4^ that can specifically detect iodide and is insensitive to chloride concentration. For HMT screening in 96-well plates, the iodide detection reagent consisted of 1 μL purified chloroperoxidase from *Curvularia inaequalis* (CiVCPO, 0.75 mg/mL) and 79 μL 3,3′,5,5′-tetramethylbenzidine (TMB, Sigma #T0565), to which 20 μL of HMT reaction I (see below) was added. For enzyme kinetics of purified HMT proteins, the iodide detection reagent contained 47 μL TMB, 0.5 μL CiVCPO (0.75 mg/mL), and 2.5 μL HMT reaction I. For the standard curve (Extended Data Fig. 13), 2.5 μL of potassium iodide (KI) at varying concentrations (5 μM to 500 μM) was added to 47 μL TMB and 0.5 μL CiVCPO (0.75 mg/mL), and absorbance at 570 nm was measured (Extended Data Fig. 10) using a Tecan Infinite plate reader.

### High throughput screening for ethyltransferase activity of HMT

Single colonies of HMT mutants in BL21 (Δmtn) cells were picked and inoculated into 800 μL LB broth supplemented with 50 μg/mL kanamycin for preculture. From preculture, 300 μL was used for glycerol stocks and stored at −80 °C. Then, 50 μL of the preculture was inoculated into 1 mL of TB (96-deepwell plate, 50 μg/mL kanamycin, 100 μM isopropyl β-D-1-thiogalactopyranoside (IPTG)) and incubated overnight at 30 °C, 900 rpm in Cytomat automated shaking incubator. Cells were pelleted at 2900xg for 15 minutes, then lysed in 300 μL of lysis buffer (1.5 mg/mL lysozyme, 0.1x Bugbuster reagent, 10 μg/mL DNase I, 1x Halt Protease inhibitor, 50 mM sodium phosphate buffer, pH 7.5). Lysis occurred at 30 °C for 1.5 hours at 900 rpm, followed by centrifugation for 20 minutes at 2900xg at room temperature.

For Reaction I, 160 μL of crude cell lysate was mixed with 20 μL SAH (1 mM) and 20 μL ethyl iodide (5 mM), both freshly prepared in DMSO. After one hour incubation at room temperature, 20 μL of Reaction I was added to 80 μL of Reaction II (1 μL CiVCPO + 79 μL TMB), and absorbance at 570 nm was measured using Tecan Infinite plate reader. A Python script calculated the slope of the increase in absorbance for each well, then calculated the fold change relative to the wild type. Each screening plate contained 90 new mutants with controls (two wells each of wild type, V140T, and empty vector). For screening V140T/S99T triple mutants, each 96-well plate contained duplicates of 36 triple mutants containing V140T/S99T, six triple mutants from the third round, and the same controls as above.

### HMT kinetics assay

The Michaelis-Menten kinetics for HMT and its mutants were determined for ethyl iodide and methyl iodide. A 50 μL reaction was prepared with 1 mM SAH, 10 μL haloalkane, and purified HMT in 50 mM sodium phosphate buffer (pH 7.5). To prevent hydrolysis, 5x haloalkane stocks were freshly prepared in DMSO. For ethyl iodide, purified wild-type HMT (wtHMT) was used at a final concentration of 87 μM (2.4 mg/mL), V140T, round 2, and round 3 mutants at 29 μM, and round 4 mutants at 8.7 μM. For methyl iodide, wtHMT was used at 0.145 nM, and V140T, round 2, round 3, and round 4 mutants at 0.29 nM. After a 10-minute incubation, 2.5 μL of the above reaction was added to the iodide assay reagent (0.5 μL CiVCPO + 47 μL TMB), and absorbance at 570 nm was measured to determine iodide concentration. For the kinetic assay, the concentration of haloalkanes was varied while keeping HMT and SAH concentration constant. The specific activity was calculated at 15 mM ethyl iodide for all purified HMT proteins. Autohydrolysis rates for each haloalkane concentration were simultaneously measured and subtracted from enzymatic reaction rates. The amount of iodide produced per minute was calculated using the standard curve (Figure S9) and fit to the Michaelis-Menten model using Origin software. All reactions were performed in triplicate.

### Purification of HMT proteins

We followed the published protocol^4^ for purifying chloroperoxidase from *Curvularia inaequalis* (CiVCPO) and stored in 50 mM sodium phosphate buffer (pH 8.0) with 100 µM sodium orthovanadate. For the iodide detection assay, 0.375 mg CiVCPO was added per 50 μL reaction.

For HMT purification, starter cultures were prepared from glycerol stocks of mutant libraries in BL21 Δmtn (DE3) strain. For protein expression, a 1:100 preculture was added to TB (50 μg/mL kanamycin) and grown at 37 °C until the OD600 reached ∼0.6. Cells were cooled and incubated overnight at 20 °C with 200 μM IPTG. The cells were harvested by centrifugation at 10,000xg at 4 °C for 10 minutes, and the pellet was resuspended in lysis buffer (20 mM sodium phosphate, 0.5 M sodium chloride, 20 mM imidazole, 1× PIC, pH 7.5). Cells were lysed by sonication (10 min on ice, 20% amplitude), followed by centrifugation (12,000xg, 20 min at 4 °C). His6-tagged proteins were purified using immobilized metal-affinity chromatography (IMAC). Lysates were clarified through a 0.45 μm filter, and 2 mL of pre-washed nickel beads (Nuvia™ IMAC Resin, Bio-Rad #7800800) were added. After 45 minutes of incubation and mixing at 4 °C, the lysates were centrifuged at 2900xg, and the supernatant was discarded. The affinity beads were transferred to a column and washed with 10x volume of lysis buffer. Proteins were eluted with elution buffer (20 mM sodium phosphate, 0.5 M sodium chloride, and 0.2 M imidazole, pH 7.5), then desalted using PD-10 desalting columns (Cytiva #17085101) and Amicon Ultra 10 kDa centrifugal filters. The proteins were stored in 50 mM sodium phosphate (pH 7.5), and concentrations were determined using absorbance at 280 nm with a Nanodrop, using the HMT extinction coefficient (42,065 M⁻¹cm⁻¹). Purity was analyzed via SDS-PAGE (Extended Data Fig. 3).

### High throughput screening for phytase activity using 4-MUP assay

Single colonies of phytase plasmids in BL21 cells were picked and inoculated into 800 μL LB broth supplemented with 50 μg/mL kanamycin for preculture in a 96-deepwell plate. 1 mL of autoinduction media (15 g/L peptone, 30 g/L yeast extract, 6.25 mL/L glycerol, 90 mL 1 M potassium phosphate buffer pH 7, 10 mL glucose (50 g/L), 100 mL lactose (20 g/L), and 50 μg/mL kanamycin) was inoculated with 5 μL aliquot of the preculture in a 96-deepwell plate. Phytase was expressed overnight (16 hours, 37 °C, 900 rpm) in a Cytomat automated shaking incubator. Cells were pelleted by centrifugation (2900xg, 15 min, room temperature) and resuspended in 200 μL lysis buffer (1 mg/mL lysozyme, 50 mM Tris-HCl buffer, pH 7.5). Lysis was carried out by incubating at 37 °C, 900 rpm for one hour, followed by centrifugation to remove cell debris.

Phytase activity was measured using a fluorescence-based 4-MUP assay^6,37^. Cleared cell lysate (10 μL) was added to 90 μL of 1.11 mM 4-MUP (4-methylumbelliferyl phosphate, Sigma #M8168) in Tris-maleate buffer (0.2 M, pH 6.6) in black, clear-bottom plates. The increase in fluorescence (*λ*_ex_ = 354 nm / *λ*_em_ = 465 nm) was measured over time using a Tecan Infinite M1000 plate reader. Each screening plate contained 90 new mutants and six control wells (wild type, M16 mutant, empty vector, and the best mutant from the previous round). The slope of the fluorescence increase was calculated for each well using a Python script, with empty vector values subtracted at each time point for normalization.

### Standard curve for 4-MUP assay

4-Methylumbelliferone (4-MU, Sigma #M1381) was dissolved into methanol and 50 µL of dissolved 4-MU solution was mixed with 50 µL of appropriate pH buffer. For pH 4.5, the buffer used was 0.25 M sodium acetate, 1 mM calcium chloride, 0.01% Tween-20. For pH 5.6 and 6.6, 0.2 M tris maleate buffer was used. Fluorescence (*λ*_ex_ = 354 nm / *λ*_em_ = 465 nm) was measured using Tecan Infinite plate reader and plotted against 4-MU concentrations to generate standard curves for each pH (Extended Data Fig. 19).

### Phytase kinetics assay

Kinetics of phytase variants was determined using a 4-methylumbelliferyl phosphate (4-MUP) assay as previously described with some modifications (ref: 10.1007/s00253-018-9308-7). Assays at pH 5.6 and 6.6 were performed in 0.2 M tris maleate buffer while assays at pH 4.5 were performed in 0.25 M sodium acetate, 1 mM calcium chloride, 0.01% Tween-20 buffer. Standard curves for pH 4.5, 5.6 and 6.6 were prepared for different concentrations of 4-MU (4-methylumbelliferone), the fluorescent product of 4-MUP (pH 4.5: 2.5-640 μM; pH 5.6: 2.5-320 μM; pH 6.6: 3.125-40 μM) and can be viewed in Extended Data Fig. 15. 50 μL of 4-MU dissolved in methanol was mixed with 50 μL of the specified buffer followed by fluorescent measurement with *λ*_ex_ 354 nm and *λ*_em_ 465 nm on a Tecan INFINITE plate reader. To prepare enzymes for kinetic assays, proteins were first mixed with 1 mM AEBSF (4-benzenesulfonyl fluoride hydrochloride) and incubated at 37 °C for 30 minutes. 90 μL of 4-MUP substrate in respective buffer was then mixed with 10 μL of the enzyme solution followed by kinetic measurement on the TECAN infinite. Fluorescence data was converted to mM 4-MU using the appropriate standard curves then divided by the enzyme concentration and used to calculate initial reaction rates. As *Yersinia mollaretii* phytase is a tetrameric enzyme with allosteric interactions, initial reaction rates were fit using OriginLab graphing software to the Hill equation *y* = *V*_max_ * *x* / (*K*_half_ + *x*) where y is the initial reaction rate, *V_max_* is the maximum reaction rate, *x* is the substrate concentration, and *K*_half_ is the substrate concentration that enables half the maximum reaction rate.

### Purification of phytase proteins

For phytase protein purification, BL21 glycerol stock from each round was used to inoculate a starter culture in LB medium followed by growth at 37 °C for 12-16 hours. 5 mL of starter cultures were added to flasks containing 500 mL Terrific Broth (6 g tryptone, 12 g yeast extract, 2 mL glycerol, 0.085 mol potassium dihydrogen phosphate, 0.36 mol dipotassium hydrogen phosphate) supplemented with 50 μg/mL kanamycin. Flasks were then shaken at 200 rpm and 37 °C for 12-16 hours. Following growth, protein expression was induced by adding 100 µM IPTG and culturing for an additional four hours at 200 rpm and 37 °C. The cells were harvested by centrifugation at 10,000xg at 4 °C for 10 minutes, and the pellet was resuspended in lysis buffer (20 mM sodium phosphate, 0.5 M sodium chloride, 30 mM imidazole, 1× Protease Inhibitor Cocktail (Sigma #P8849), pH 7.5) supplemented with 1 mg/mL lysozyme. Cells were then lysed for 1 hour at 37°C followed by centrifugation (12,000xg, 20 min at 4°C). His6-tagged proteins were purified using immobilized metal-affinity chromatography (IMAC). Lysates were clarified through a 0.45 μm filter, and 3 mL of pre-washed nickel beads (Nuvia™ IMAC Resin, Bio-Rad #7800800) were added. After 45 minutes of incubation and mixing at room temperature, the lysates were centrifuged at 2900xg, and the supernatant was discarded. The affinity beads were transferred to a column and washed with 10x volume of lysis buffer. Protein was eluted by adding 10 mL elution buffer (20 mM sodium phosphate, 0.5 M sodium chloride, and 0.2 M imidazole, pH 7.5). Protein was concentrated and the buffer was exchanged to 0.25 M sodium acetate, pH 5.5, 1 mM calcium chloride, 0.01% Tween-20 using Amicon Ultra 10 kDa centrifugal filters. Concentrations were determined using absorbance at 280 nm with a Nanodrop and enzymes were stored at 4°C

### PAGE gels for purified proteins

For SDS-PAGE, the purified proteins were mixed with 1x Laemmli sample buffer (Biorad #1610737) with 5% 2-mercaptoethanol (Sigma #M6250) and boiled at 95°C for five minutes. An equal amount of each sample was then loaded onto a 10% precast gel (Biorad #4561034) with protein ladder (Biorad # 1610374) and run using Tris/Glycine/SDS running buffer (Biorad # 1610732). After Coomassie staining, the gels were thoroughly washed before imaging. For native gel for phytase proteins, the purified proteins were mixed with native sample buffer (Biorad #1610738) and equal amount was loaded onto 7.5% gel (Biorad #4568024) and run with Tris/glycine running buffer (Biorad # 1610771). NativeMark unstained protein standard (Invitrogen # LC0725) ladder was used for native gel.

### Worklist generation

The seamless integration of various modules required generating worklists for PCR, which were created using Python scripts. Each round of mutagenesis added one additional mutation to plasmids generated in the previous round, such that the variants in the third round of evolution contained three total mutations. Thus, linearized plasmids containing mutations from one round of screening serve as templates for the next round. A Python script efficiently selected the minimal number of PCR templates needed for the next cycle. The linearized PCR templates were transferred to a 384-well plate, and a worklist was generated to distribute them into a 96-well PCR plate. Another worklist was created to distribute the primers using Tecan Fluent in 96-well PCR plates. Primers were ordered in a 96-well format from IDT and diluted to 2 μM stocks with 12.5 μL being added to each PCR reaction. Worklists were also used for spreading the transformed *E. coli* cells in 8-well omnitray LB plates.

## Data availability

The raw data for all figures and tables are included as supplementary data files. The plasmids created during this research are available upon request.

## Code availability

The source code for zero-shot predictions and supervised learning model is available at https://github.com/Zhao-Group/closed-loop. The python scripts used for primer design and worklist generation are available at https://github.com/Zhao-Group/Primer_Design_and_Worklists.

## Acknowledgements

We thank members of the Biosystems Design theme at the Carl R. Woese Institute for Genomic Biology for their discussions and feedback. We thank Prof. Uwe Bornscheuer (University of Greifswald) for sharing research materials necessary for the *At*HMT enzyme and Prof. Ulrich Schwaneberg and Dr. Anna Joelle Ruff for helpful discussions related to the *Ym*Phytase enzyme. We also thank Nicole Setiawan for general laboratory duties that helped this project.

## Author contributions

T.Y. predicted enzyme mutations using the unsupervised and supervised models and designed the natural language agent. N.S. developed the site-directed mutagenesis protocol and designed the automated workflow while N.S. and S.L. performed automation implementation and operation. N.S. and S.L performed the four cycles of engineering of the *At*HMT and *Ym*Phytase enzymes. H.C. validated the phytase 4-MUP assay. A.R. helped with performing protein engineering experiments and SDM. N.S., S.L., and J.L. purified the *At*HMT and *Ym*Phytase protein variants and performed kinetic characterization. N.S., S.L, T.Y, and H.Z. wrote the manuscript. All authors contributed to editing the manuscript. H.Z. directed research activities and procured the funding to support this study.

## Additional information

### Data and materials availability

Correspondence and requests for materials should be addressed to Huimin Zhao.

**Supplementary information** is available for this paper.

## Extended Data Figures

**Extended Data Fig. 1.**
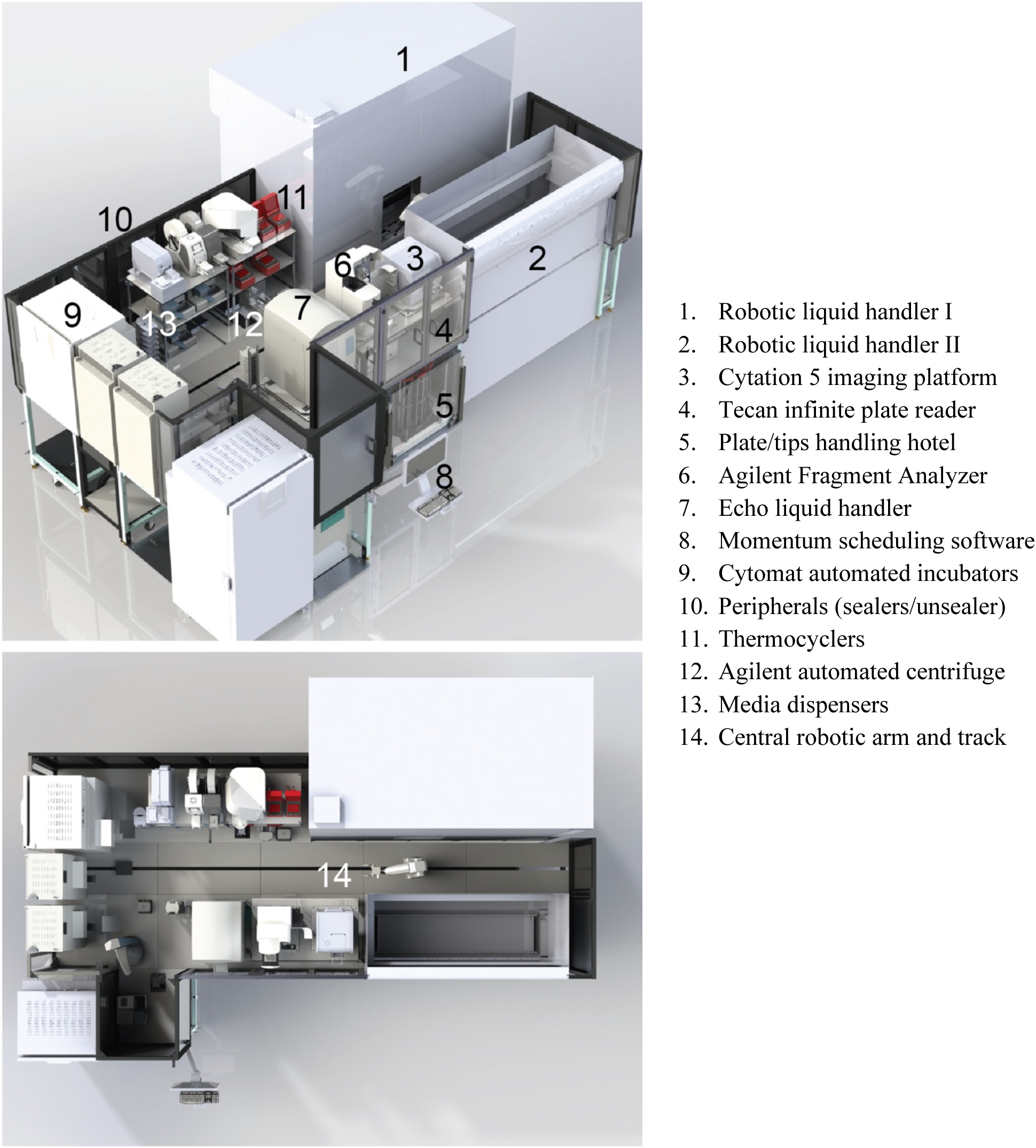
An overview of the iBioFAB automation platform. Isometric and top-down views of the iBioFAB automation platform are shown. Various core and peripheral instruments are integrated using Thermo Fisher Momentum scheduling software and a Thermo Fisher F5 central robotic arm. A description of the component instruments is listed alongside the image.

**Extended Data Fig. 2.**
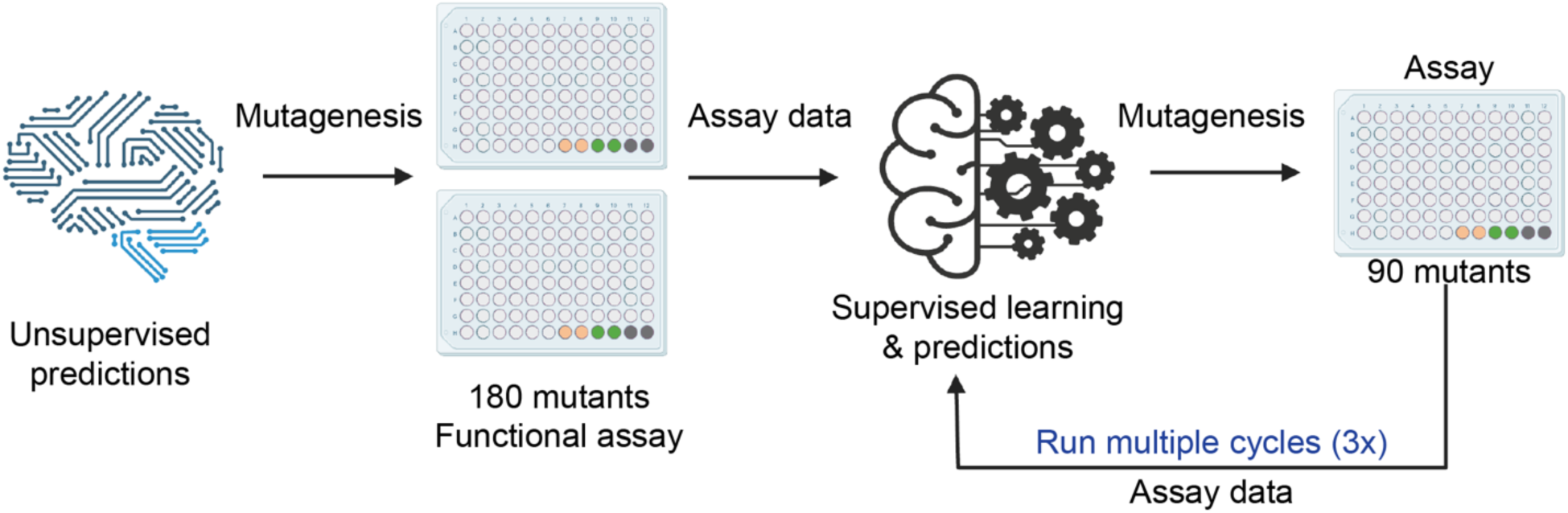
Overview of the autonomous protein engineering workflow. The workflow is initiated by predicting 180 mutants for the first round of screening in two 96-well plates. The data from each screening cycle is used to train a supervised learning model, which then predicts mutants for the subsequent cycle. After the first cycle, subsequent screening cycles used a single 96-well plate, containing 90 new mutants and six controls (two empty vectors, two wild type enzymes, and two positive control enzymes).

**Extended Data Fig. 3.**
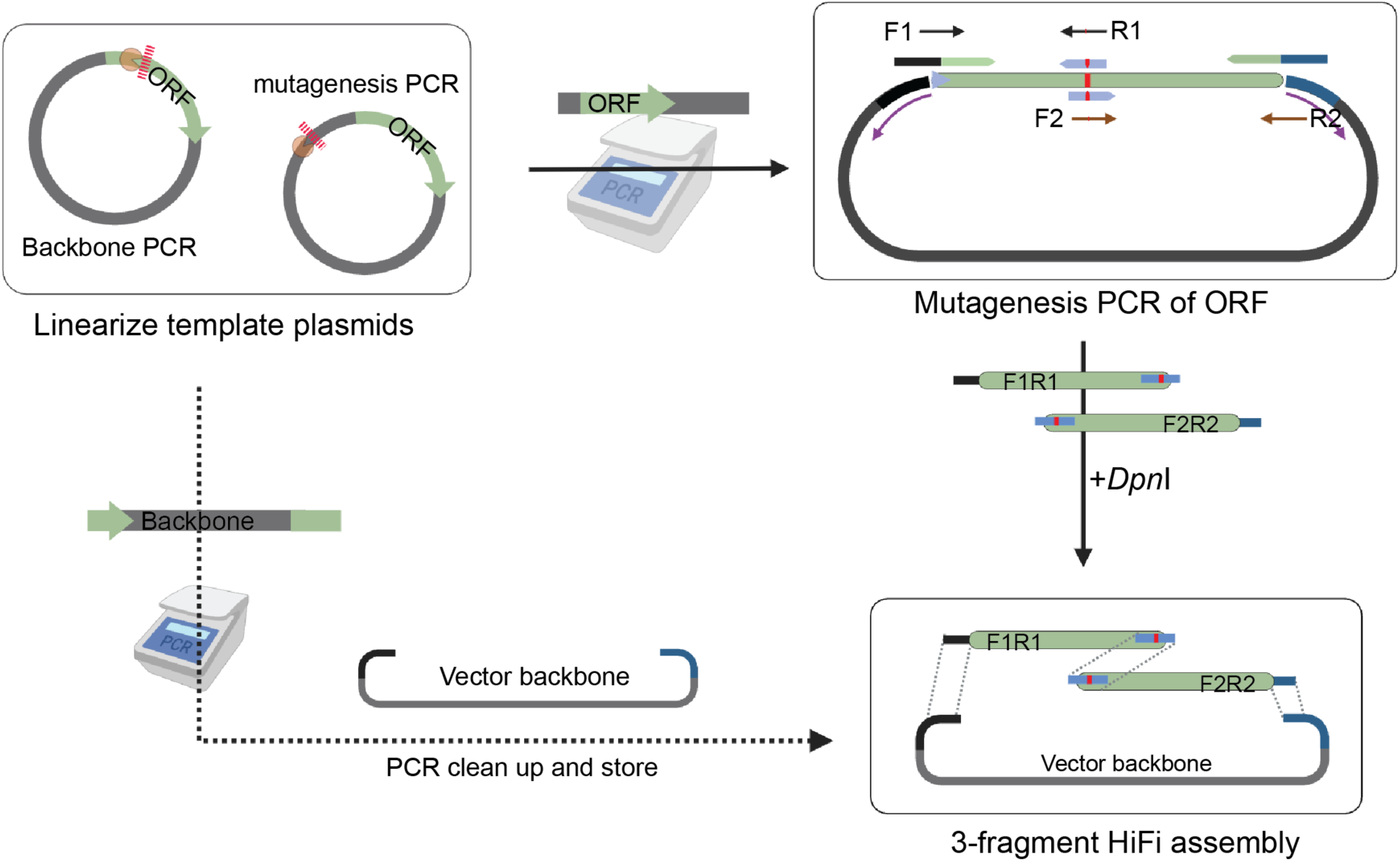
Detailed schematics for the site-directed mutagenesis protocol. The site-directed mutagenesis protocol used a 27 bp mutagenic primer (12+3+12) for generating mutations in the ORF. Each mutation site required two mutagenic primers to amplify the adjacent fragments. Another set of primers were used for PCR of vector backbone. The three PCR fragments contained overlapping regions of 27–40 bp. The template plasmids for PCRs were linearized with appropriate restriction enzyme. For ORF PCR, ∼200 pg of template DNA was used per reaction. The PCR reactions were treated with *Dpn*I to eliminate the original templates, followed by HiFi assembly of the backbone and two adjacent ORF fragments.

**Extended Data Fig. 4.**
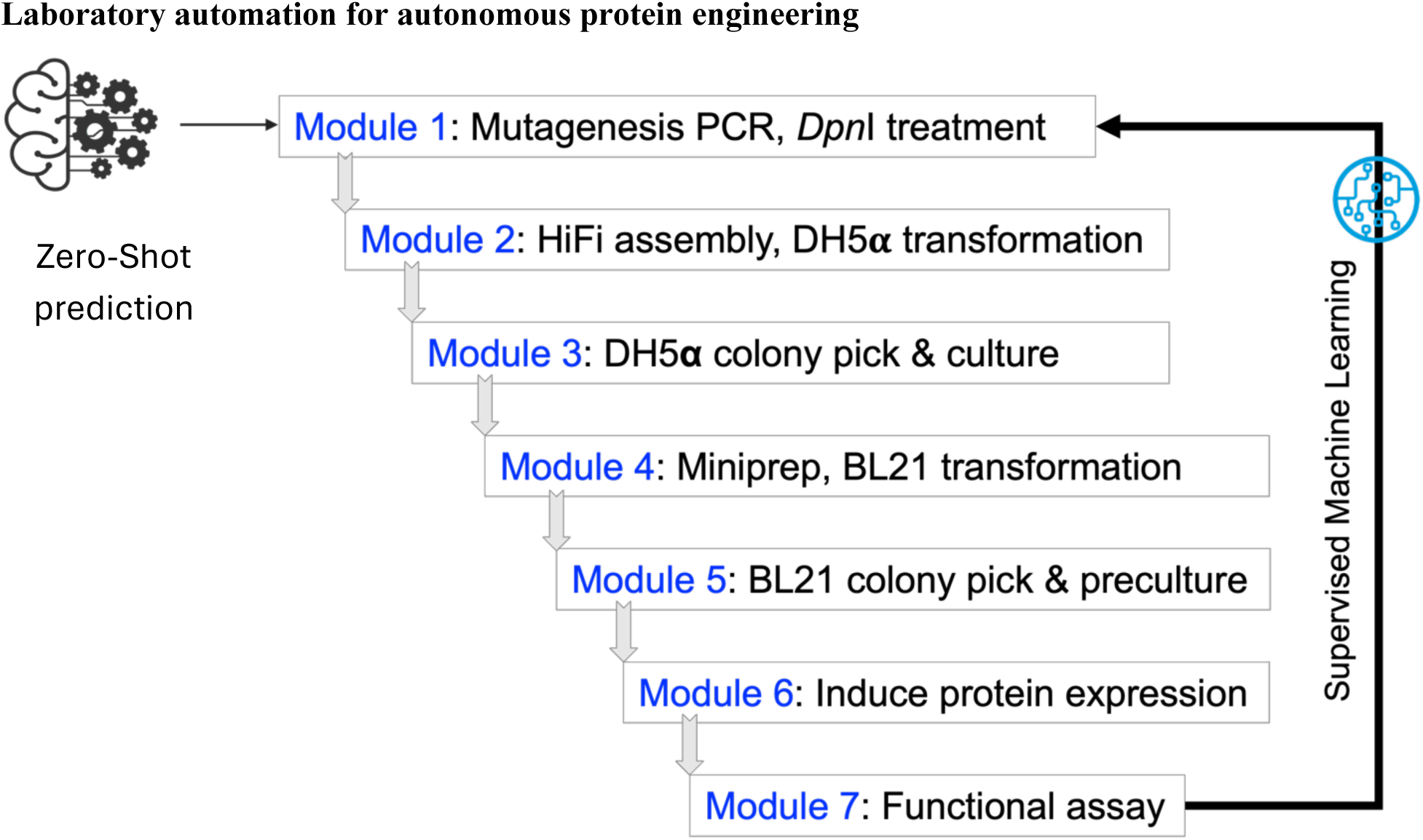
Overview of the laboratory automation workflow for autonomous protein engineering. The experimental workflow was divided into seven modules, with each designed to be completed in one day. Each module consisted of multiple experiments, coordinated and integrated using Thermo Fisher Momentum scheduling software across one or more instruments on the iBioFAB. The first two rounds of HMT screening served as a test case to design, develop, and troubleshoot the workflow, addressing various experimental and instrumental issues for a seamless, interconnected process. Detailed operations for each module are described in the subsequent extended data figures.

**Extended Data Fig. 5.**
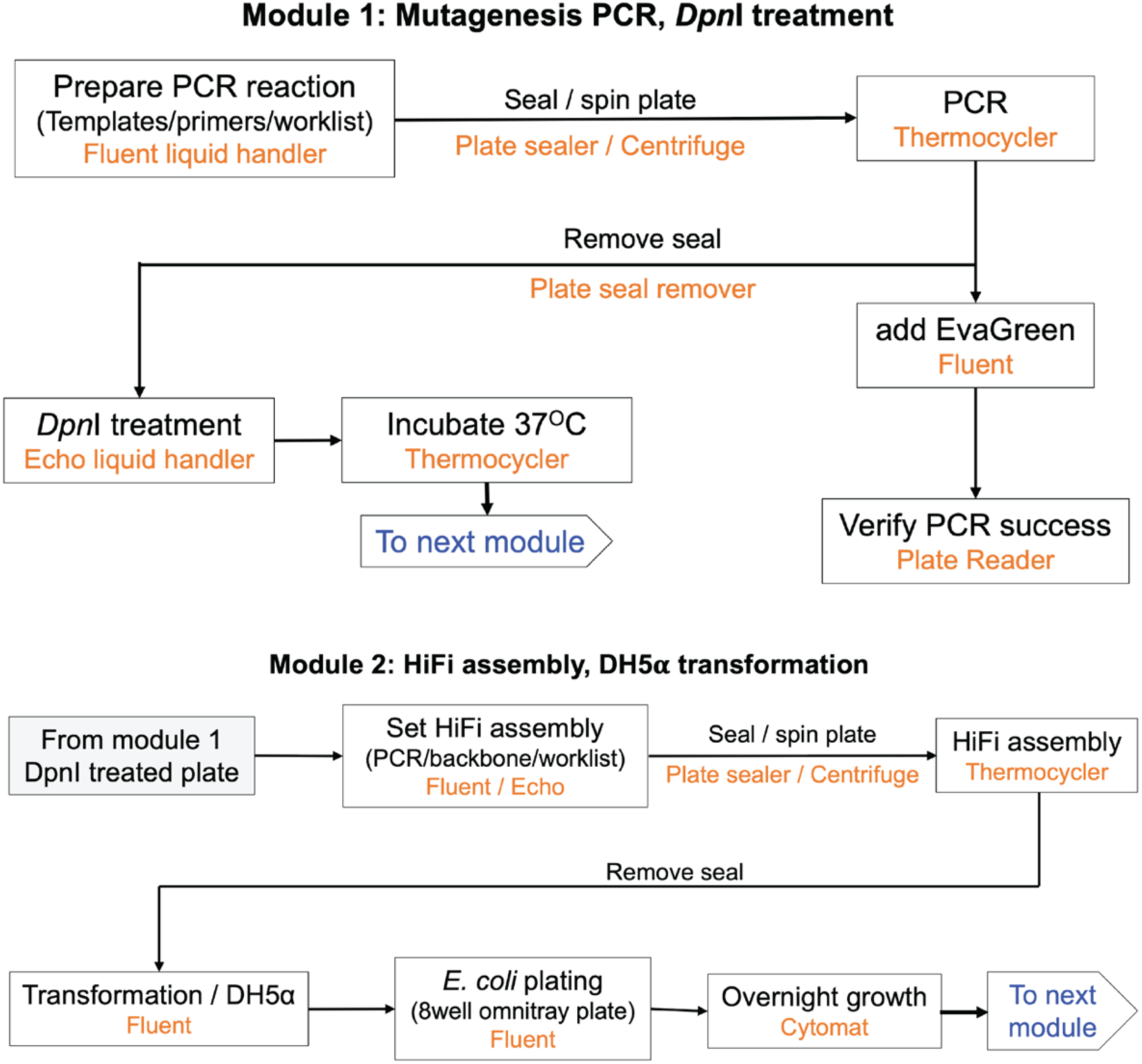
Detailed overview of modules 1 and 2 for automated protein engineering. Module 1 of the workflow prepared mutagenesis PCR for 96 mutants in a 96-well PCR plate. The templates and primers for PCR were mixed using worklists on the Tecan Fluent and Echo liquid handler. PCR success was measured by adding 2.5 μL of the PCR reaction to 47.5 μL of 1x Evagreen dye (Biotium #31000) and measuring fluorescence (*λ*_ex_ = 498 nm / *λ*_em_ = 535 nm) with the Tecan Infinite plate reader. 25 μL PCR product was transferred to a new PCR plate, and 1 μL of *Dpn*I (NEB #R0176) was added, followed by incubation at 37 °C. The *Dpn*I-treated PCR products were then transferred to a 384-well plate, along with the vector backbone, and a worklist guided the mixing of the correct fragments in the Echo liquid handler for HiFi assembly. After a 30-minute HiFi assembly at 50 °C, competent DH5α cells in a 96-well plate were transformed by heat shock on the Tecan Fluent using onboard heating/cooling blocks. The cells were plated on 8-well omnitray agar plates containing LB + 50 μg/mL kanamycin and incubated overnight at 37 °C in a Cytomat automated shaking incubator.

**Extended Data Fig. 6.**
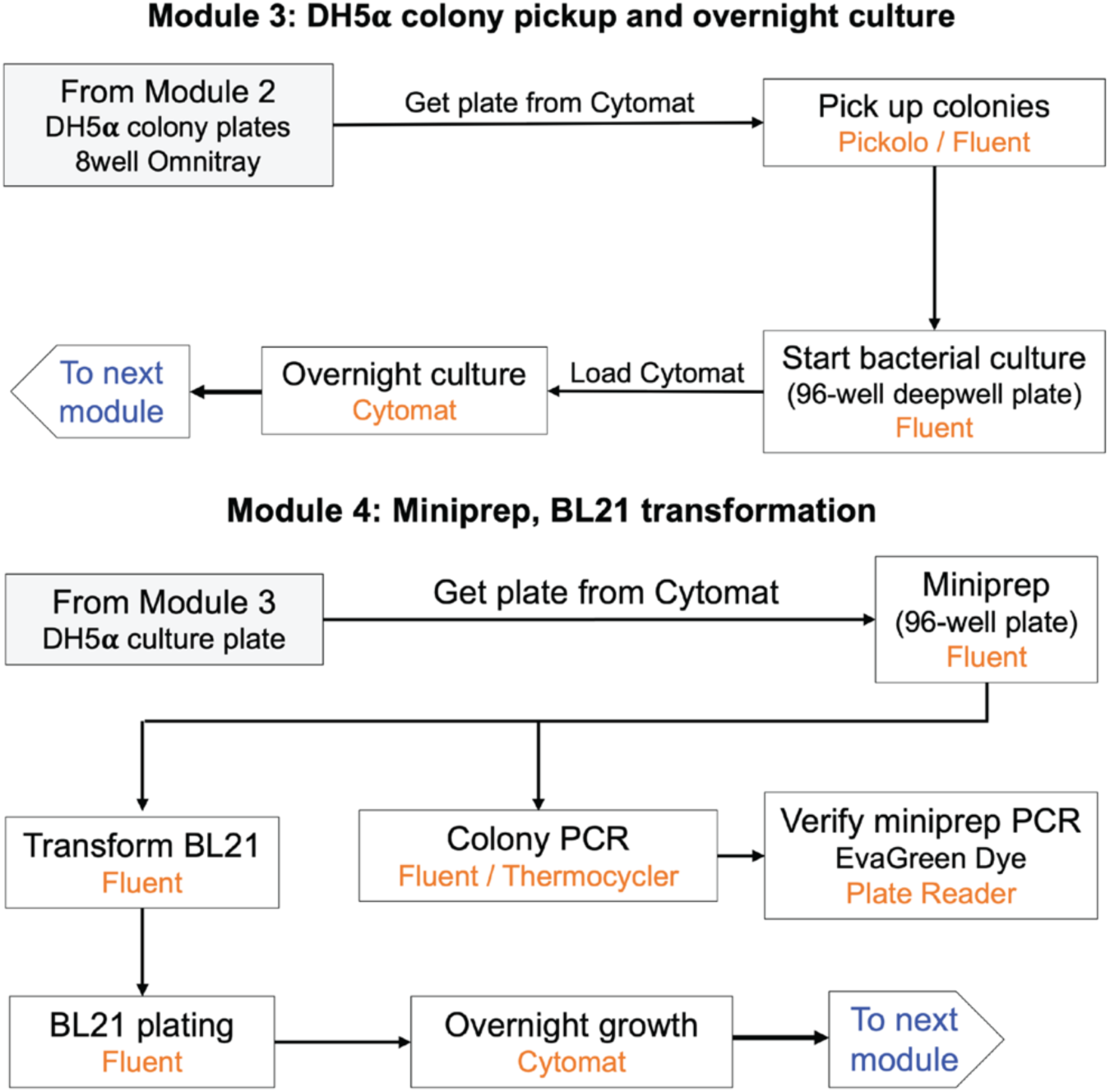
Detailed overview of modules 3 and 4 of laboratory automation workflow. In Module 3, the Pickolo (SciRobotics) system on the Tecan Fluent was used to pick DH5α colonies from 8-well omnitray plates and inoculate them into 1 mL of TB media (kanamycin 50 μg/mL) in a 96-deepwell plate. The inoculation process maintained the order of mutations in the 96-well plate to match the initial placements in the PCR plate. After overnight culture in an automated Cytomat incubator at 37 °C and 900 rpm, miniprep was performed using the vacuum module on the Tecan Fluent with the PureLink Pro Quick96 Plasmid Purification Kit (Invitrogen #K211004A). 5 μL miniprep DNA was used to transform competent BL21 cells in a 96-well plate by heat shock, using the heating/cooling block onboard the Tecan Fluent. The BL21 cells were plated on 8-well LB/agar omnitray plates (kanamycin 50 μg/mL) and incubated overnight at 37 °C in the Cytomat. We also performed a colony PCR and verified successful PCRs using Evagreen dye to measure the success of HiFi assembly. We consistently achieved a >90% success rate for HiFi assembly. Since we started with 96 mutants and only needed 90 for screening, the <10% failed HiFi reactions did not impact the workflow. The minipreps of some mutants were randomly selected and sent for sequencing. Throughout all evolutionary cycles of both HMT and phytase, we observed 100% correct mutagenesis for the randomly chosen mutants that were sent for sequencing.

**Extended Data Fig. 7.**
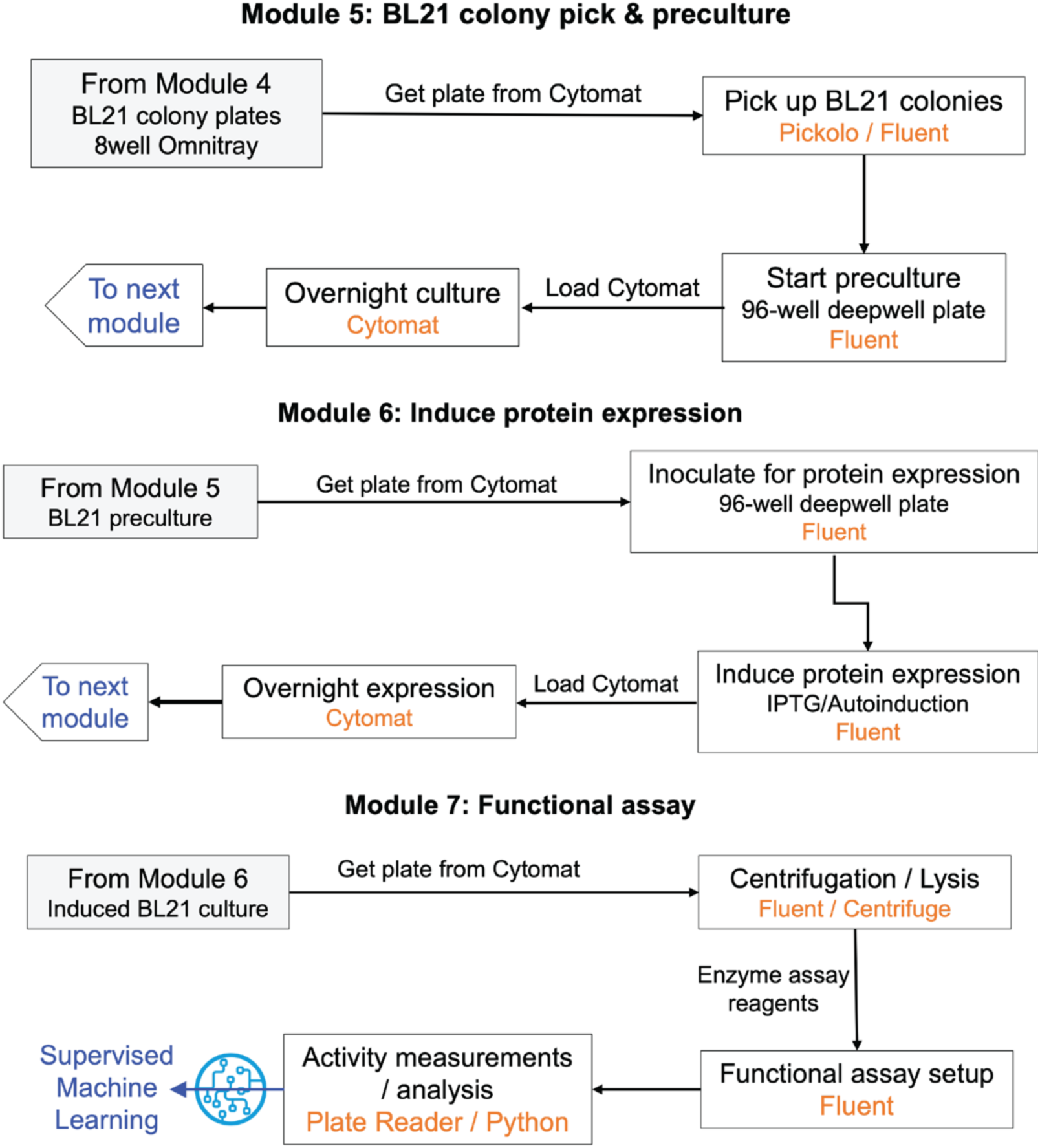
Detailed overview of modules 5, 6 and 7 of laboratory automation workflow. In Module 5, BL21 colonies were picked using the Pickolo (SciRobotics) system on the Tecan Fluent and inoculated into LB media (kanamycin 50 μg/mL) in a 96-deepwell plate for preculture. Control wells, including wild type, empty vector, and positive control, were also added and inoculated at this stage. The next day, Module 6 inoculated a 96-deepwell plate for protein expression using the preculture, and the plates were loaded into the Cytomat under the appropriate conditions for protein expression. In Module 7, the cells were centrifuged, followed by lysis, a second centrifugation, and the collection of lysates for the activity assay. The protein activity assays were optimized to be compatible with liquid handling systems. Protein activity was measured using the Tecan Fluent plate reader, and the data were analyzed using Python scripts to calculate the relative activity of mutants compared to the wild type.

**Extended Data Fig. 8.**
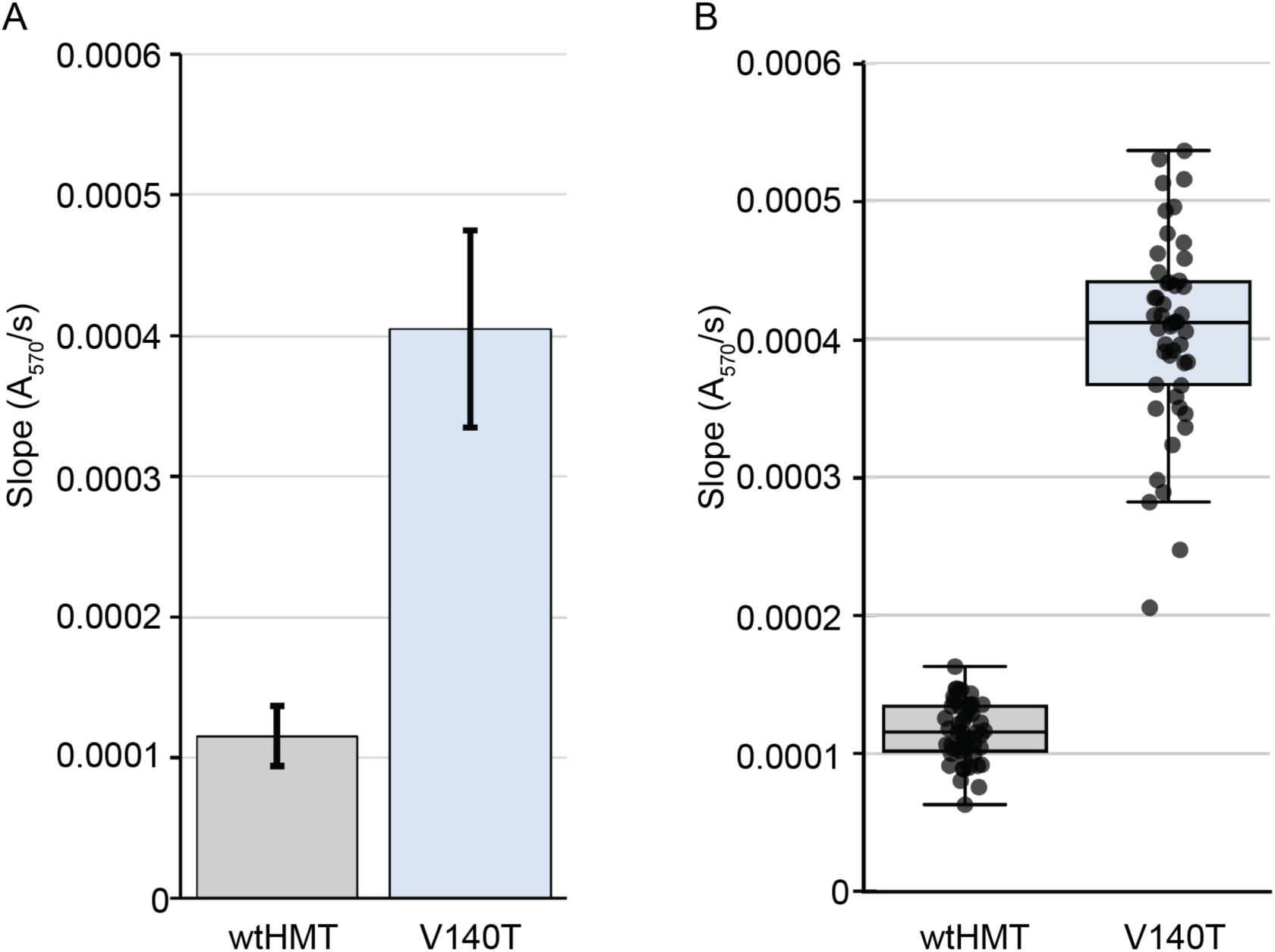
Variance of iodide detection assay for HMT activity in 96-well plate. In a 96-well plate, wtHMT and the positive control V140T mutant were each expressed in 48 wells. Subsequently, the lysates from the plate were used to detect HMT activity using the iodide detection assay. Absorbance at 570 nm was measured and the rate of increase in A_570_ was calculated. (**A**) The average and standard deviation for the wtHMT and the V140T mutant were calculated from the 48 wells, bars represent average with errors bars for standard deviation. (**B**) A box plot with overlaying individual values is shown for both the wild type and V140T.

**Extended Data Fig. 9.**
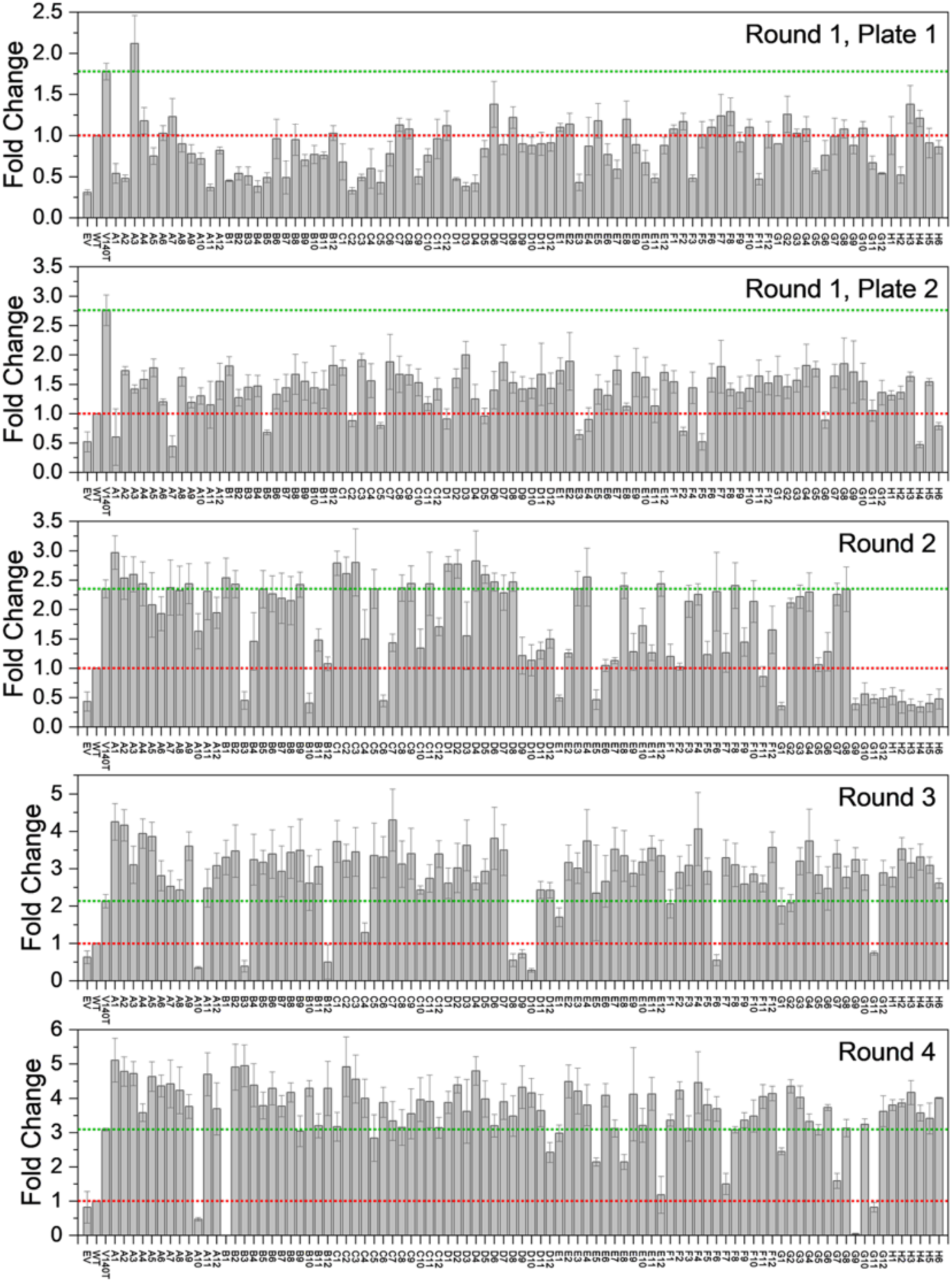
Crude lysate screening from all four rounds of autonomous protein engineering of *Arabidopsis thaliana* halide methyltransferase (*At*HMT). *At*HMT mutants were screened for activity in 96-well format using the iodide assay, and the fold change compared to the wild type was plotted. Each plate contained ∼90 new mutants. Round 1 screening involved two 96-well plates for 180 single mutants, followed by 90 mutants with double, triple, and quadruple mutations in rounds 2, 3, and 4, respectively. Each screening plate included the wild type (red dotted line), the V140T mutant (green dotted line), and an empty vector (EV). The plots represent the average of n ≥ 3 experiments with standard deviations indicated by error bars. Mutants are labeled according to the wells of the 96-well plate, and the names of the mutations are provided in Supplementary Table 3 along with the original data.

**Extended Data Fig. 10.**
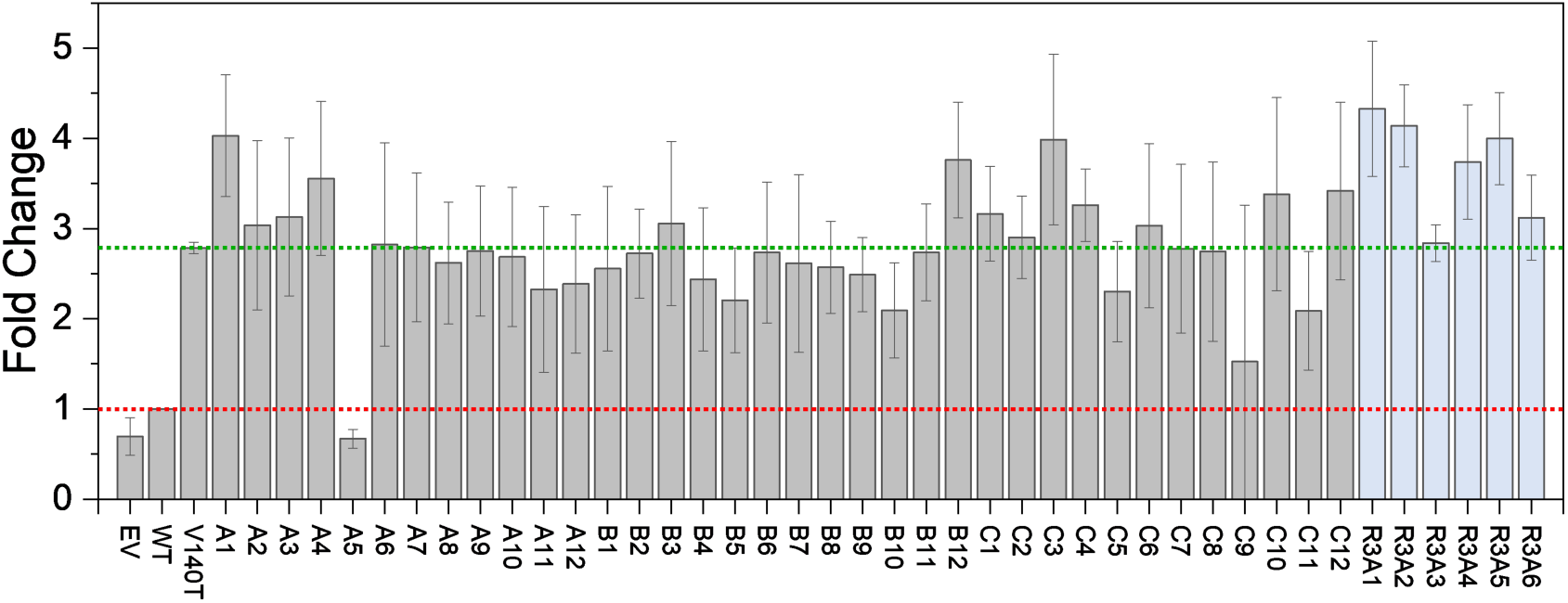
Comparing human intuition with machine learning predictions for *At*HMT variants. The predicted mutants for third round (triple mutants) did not include mutants with a V140T/S99T combination in the top 500 predictions, despite V140T/S99T being one of the best double mutants from the second evolutionary round. We generated the top 36 triple mutants containing V140T/S99T (grey bars) and screened them for *At*HMT activity, along with the top six predicted triple mutants for the third round (light blue bars). The screening plate also contained the wild type (red dotted line), V140T (green dotted line), and empty vector controls. The fold change compared to the wild type is plotted as average of four independent experiments with standard deviations indicated by error bars. The names of the mutations corresponding to each well are listed in Supplementary Table 5 along with the original data.

**Extended Data Fig. 11.**
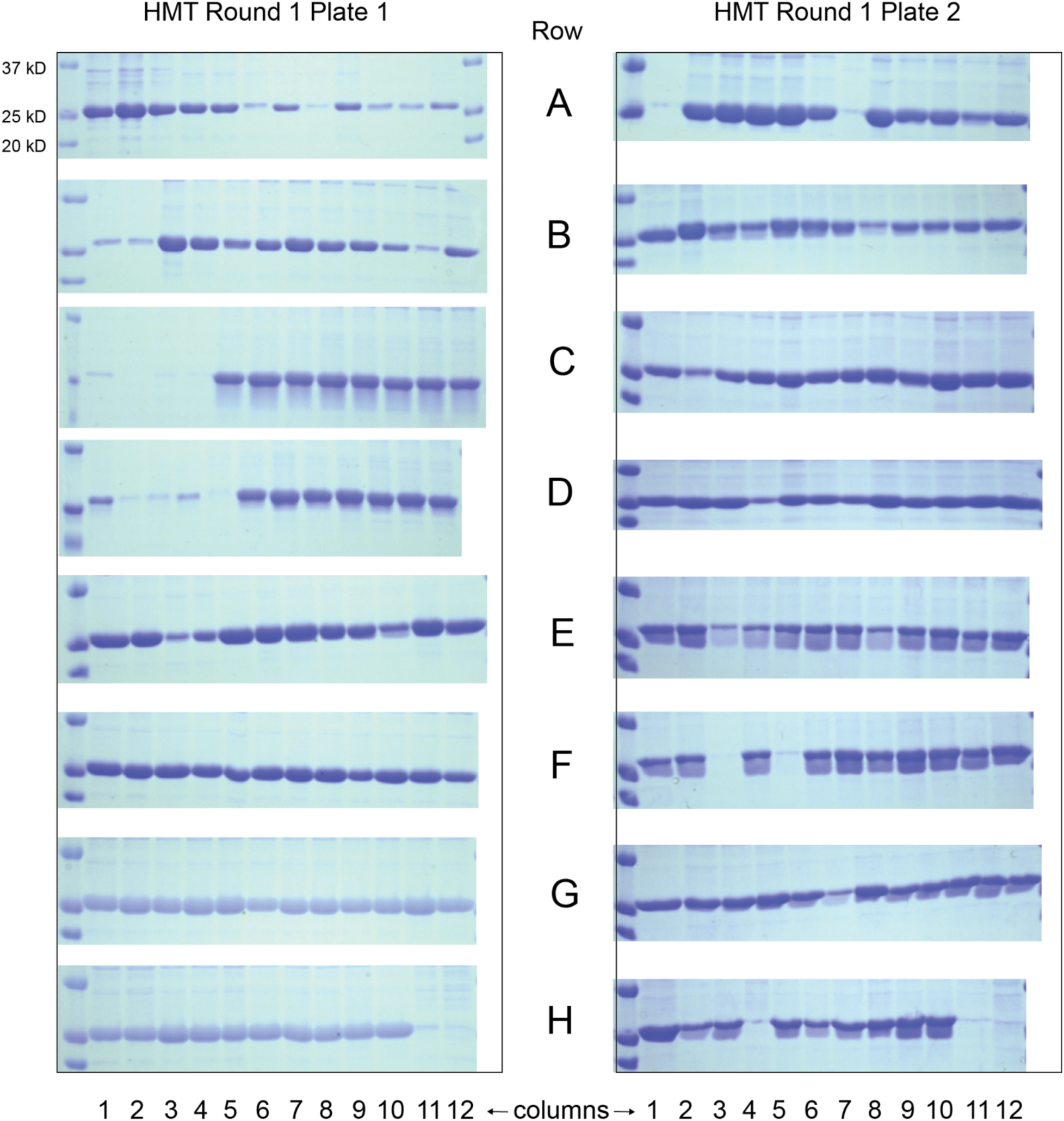
SDS-PAGE of cell lysates from plates 1 and 2 from the first round of autonomous protein engineering of *At*HMT. Equal amounts of lysates from three replicate plates were combined and mixed with 2x Laemmli buffer (+5% BME) and boiled for five minutes at 95 °C. Subsequently, 15 µL of each sample was run on an 10% SDS-PAGE gel and stained with Coomassie Blue (Bio-Rad #1610436). Images were acquired after thoroughly washing away the stain. The newly generated mutants were loaded in wells A1 to H6, wtHMT in H7 and H8, V140T in H9 and H10, and the empty vector in H11 and H12. The names of the mutations are available in Supplementary Table 3.

**Extended Data Fig. 12.**
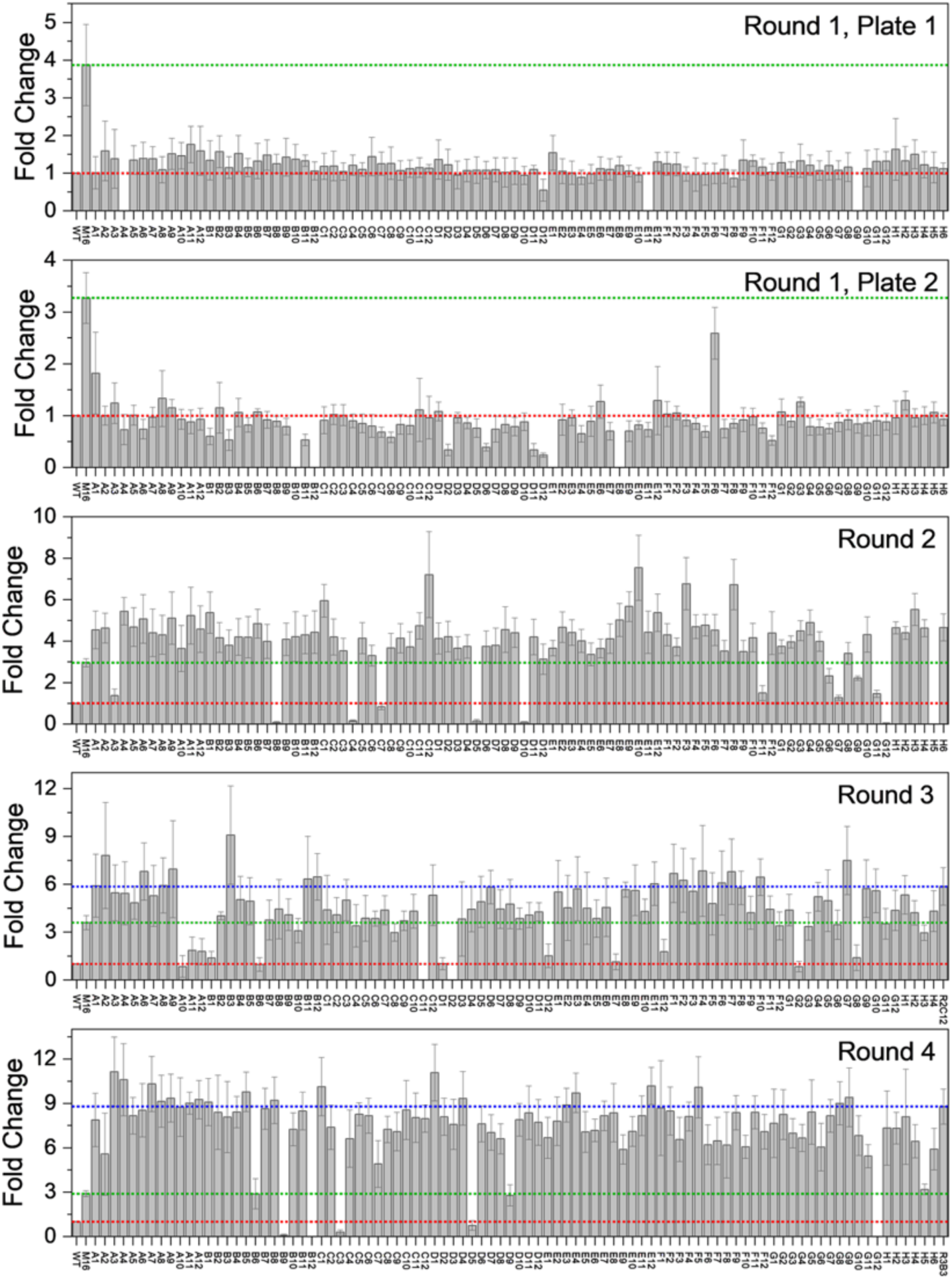
Crude lysate screening from all four rounds of autonomous protein engineering of *Yersinia mollaretii* phytase (*Ym*Phytase). *Ym*Phytase mutants were screened in 96-well format using the 4-MUP assay, and the fold change compared to the wild type was plotted. Each 96-well plate contained 90 new mutants and six controls, with the first round of evolution involving two 96-well plates for 180 single mutants and 12 controls, followed by approximately 90 mutants with double, triple, and quadruple mutations in the second, third, and fourth rounds, respectively. Each screening plate included the wild type (red dotted line), a previously reported M16 variant containing a T44V/K45E double mutation *(7)* (green dotted line), and an empty vector (EV). To calculate the slope, the fluorescence background for EV was subtracted at each time point. For the third and fourth rounds, one of the best mutants from the previous round was included (blue dotted line). The plots represent the average of n ≥ 3 experiments with standard deviation. Mutants are labeled according to the wells of the 96-well plate, and the names of the mutations are listed in Supplementary Table 4 along with the original data.

**Extended Data Fig. 13.**
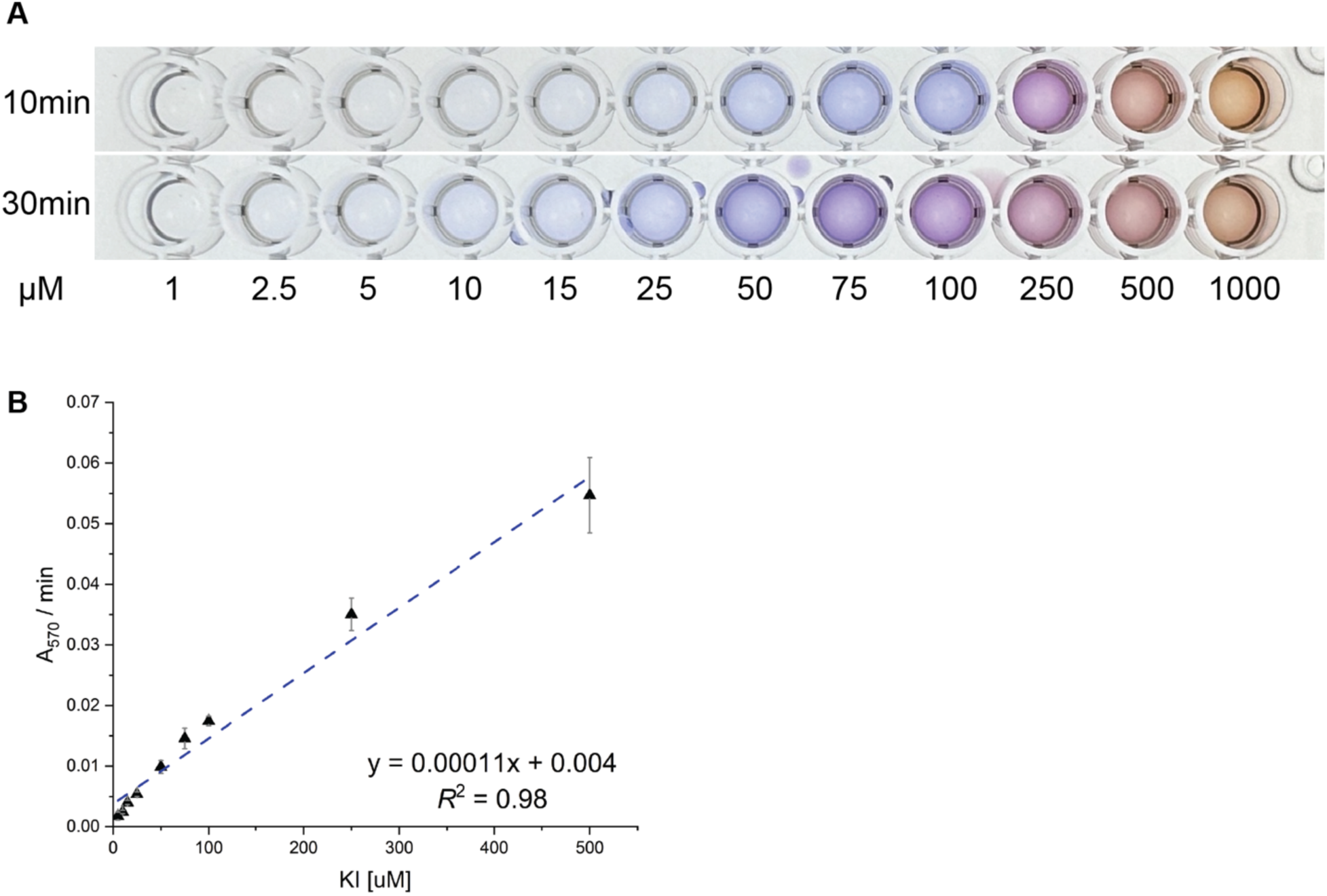
Iodide detection assay and the standard curve for iodide detection. (**A**) Image of color formation after adding different concentrations of potassium iodide (KI) to the iodide detection reagent at 10 minutes and 30 minutes (**B**) Standard curve for iodide detection assay. In a 50 μL reaction, 2.5 μL of potassium iodide (KI) [5 μM to 500 μM] was added to the iodide detection reagent containing TMB (47 μL) and CiVCPO (0.5 μL / 0.35 mg). The initial increase in Absorbance at 570 nm was measured and slope per minute was plotted against KI concentration (n=4).

**Extended Data Fig. 14.**
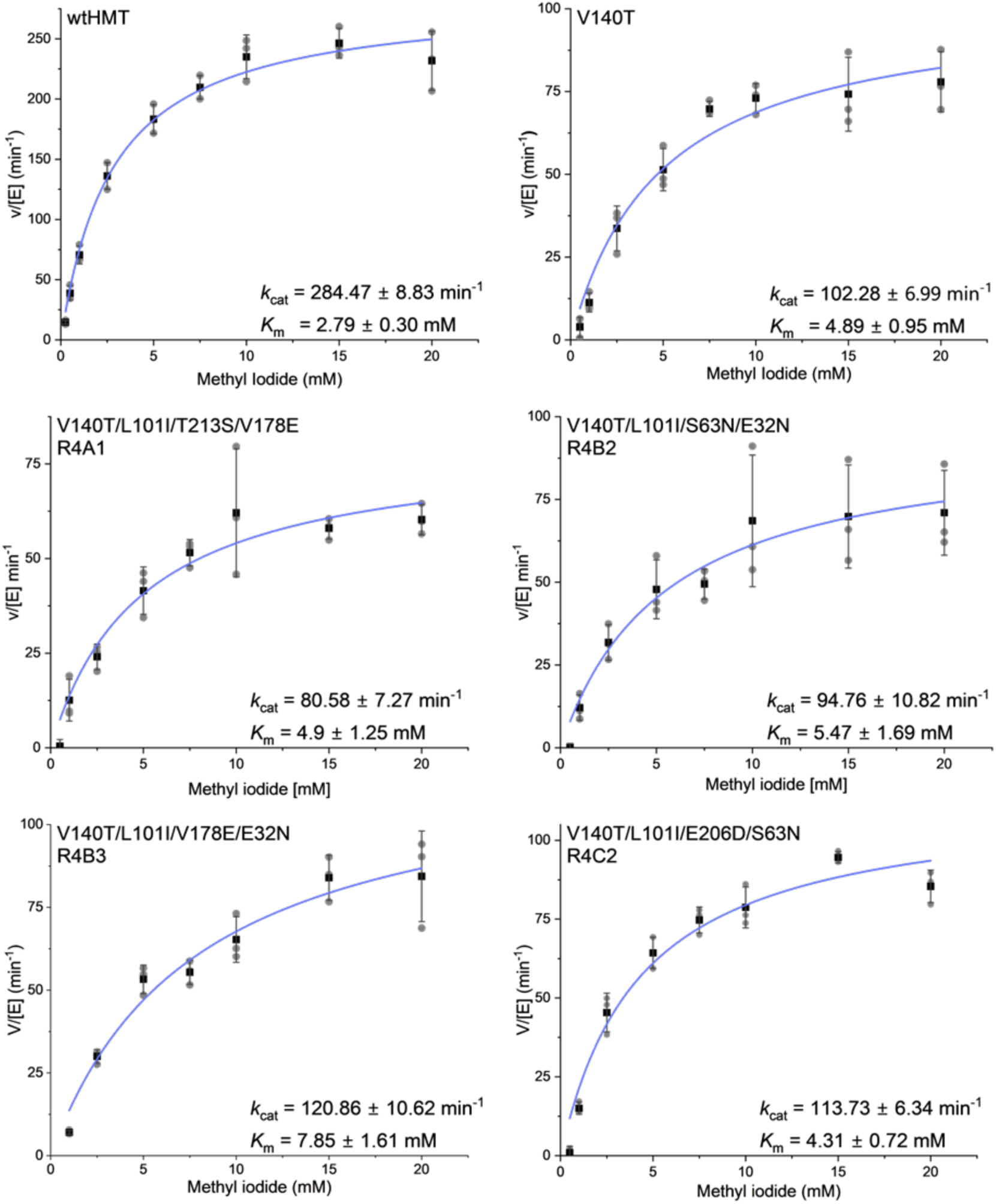
Michaelis-Menten kinetics of wild type *At*HMT and its round 4 variants using methyl iodide. The methyltransferase activity for purified variants of *At*HMT was determined using different methyl iodide concentrations. wtHMT was added at 200 ng (0.145 μM) per reaction while the round 4 variants were added at 400 ng (0.29 μM) per reaction. The curves were fit using Michaelis Menten fit on OriginPro 2023. The values (circle), mean (square), fitted plot (blue), and standard deviation are plotted (n=3). The name of the purified protein is labeled on each graph. The notation R4C2 represents well C2 of the fourth round of autonomous protein engineering.

**Extended Data Fig. 15.**
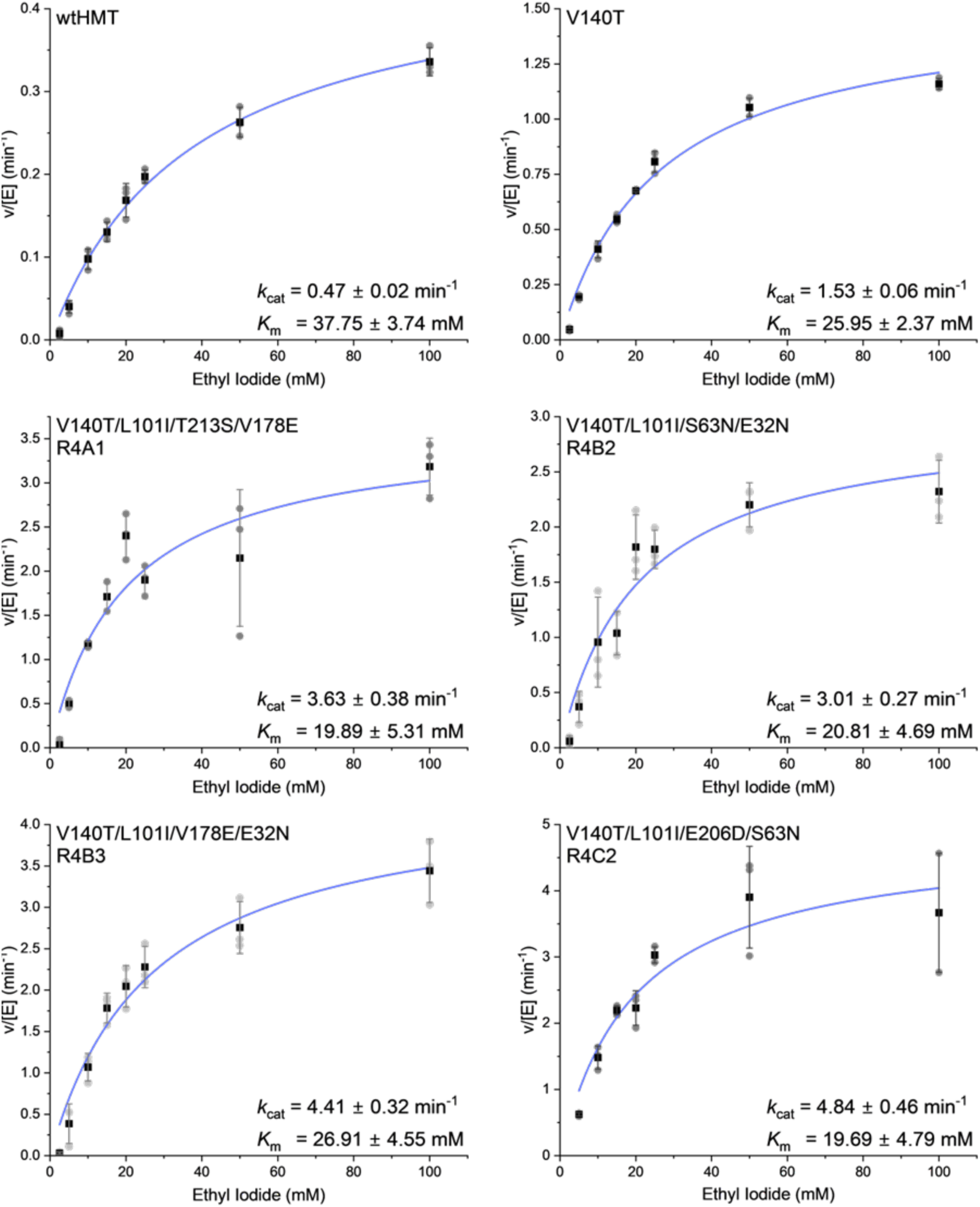
Michaelis-Menten kinetics of wild type *At*HMT and its round 4 variants using ethyl iodide. The ethyltransferase activity for purified variants of *At*HMT was determined using different ethyl iodide concentrations. wtHMT was added at 87 μM (2.4 mg/mL), V140T at 29 μM, and round 4 variants at 8.7 μM per reaction. The curves were fit using Michaelis Menten fit on OriginPro 2023. The values (circle), mean (square), fitted plot (blue), and standard deviation are plotted (n=3). The name of the purified protein is labeled on each graph. The notation R4C2 represents well C2 of the fourth round of autonomous protein engineering.

**Extended Data Fig. 16.**
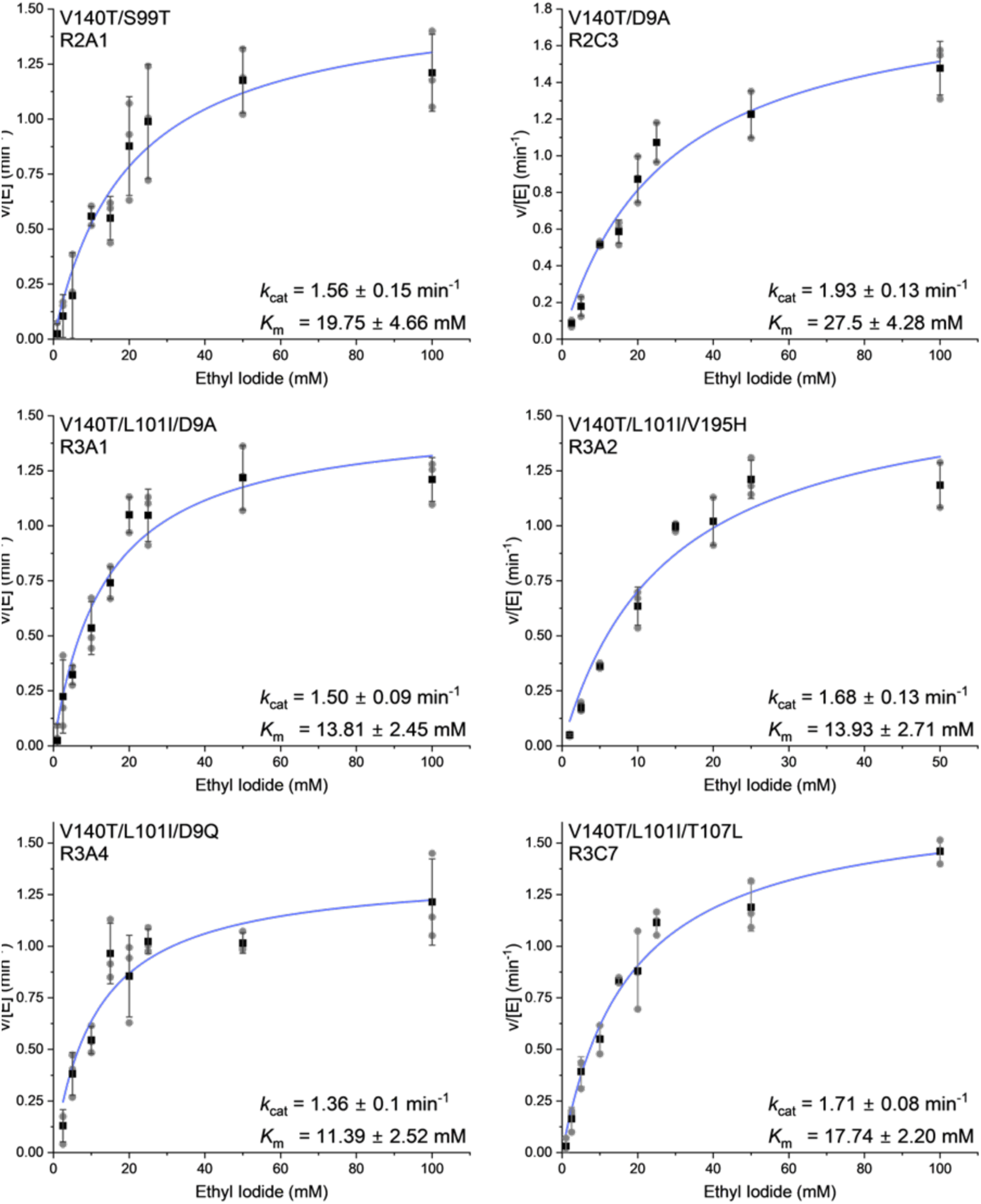
Michaelis-Menten kinetics of wild type *At*HMT and its round 2 and round 3 variants using ethyl iodide. The ethyltransferase activity for purified variants of *At*HMT was determined using different ethyl iodide concentrations. Second and third round variants were added at final concentrations of 29 μM per reaction. The curves were fit using Michaelis Menten fit on OriginPro 2023. The values (circle), mean (square), fitted plot (blue), and standard deviation are plotted (n=3). The name of the purified protein is labeled on each graph. The notation R3C7 represents well C7 of the third round of autonomous protein engineering.

**Extended Data Fig. 17.**
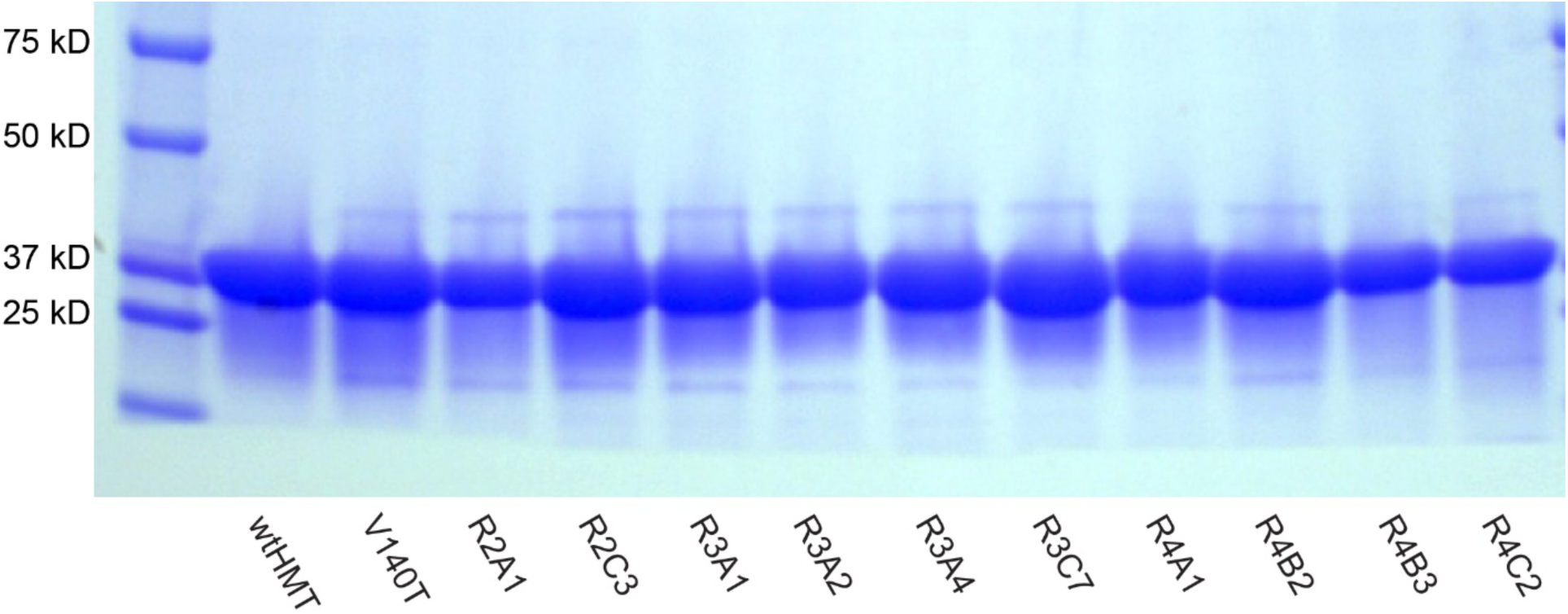
SDS PAGE of purified *At*HMT proteins. The purified proteins were mixed with 1x Laemmli buffer (+5% BME) and boiled at 95 °C for five minutes. An equal amount of each sample was loaded on a 12% SDS-PAGE gel along with a protein ladder (BioRad # 1610374). After Coomassie staining, the gels were thoroughly washed before imaging.

**Extended Data Fig. 18.**
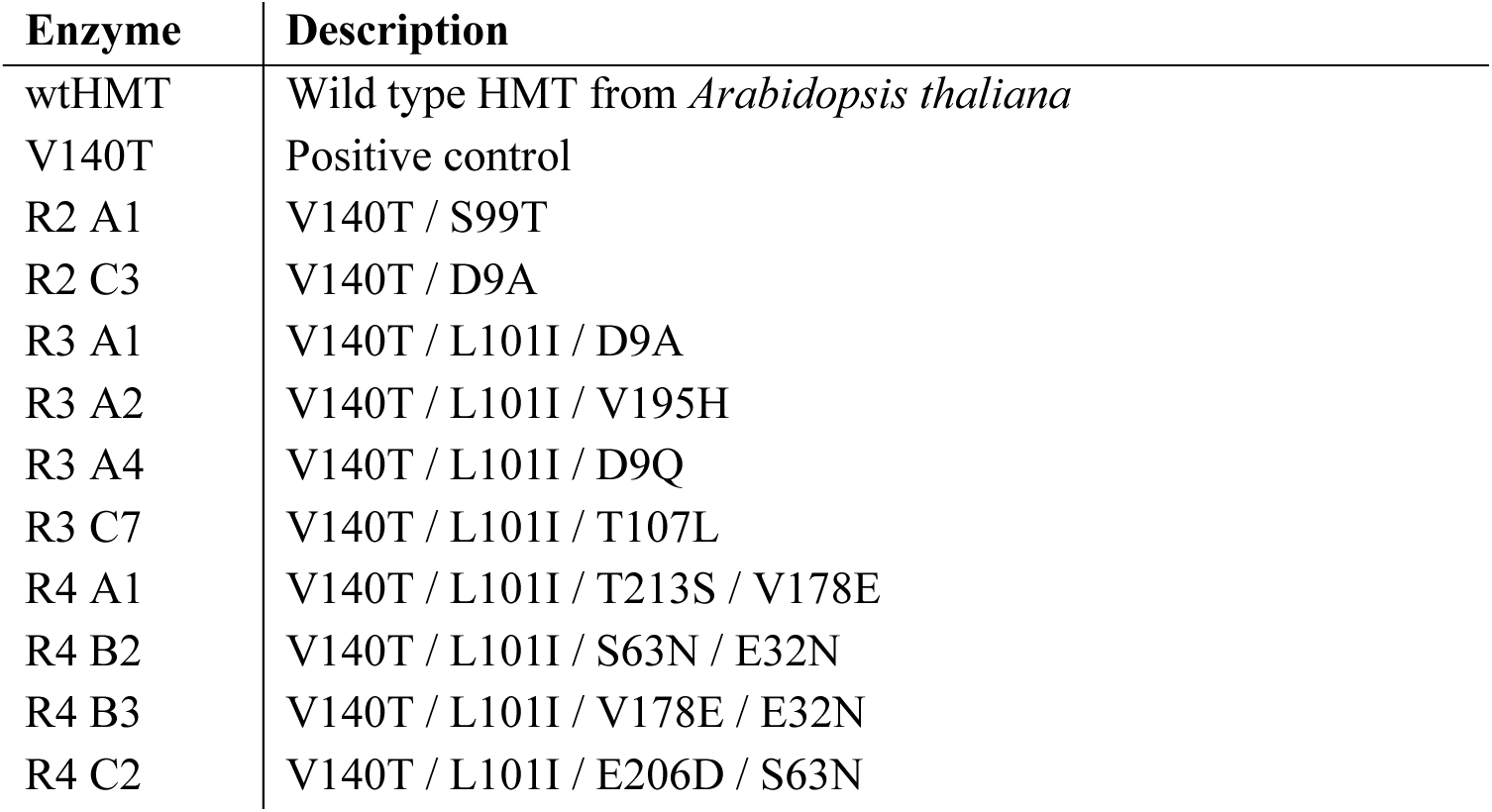
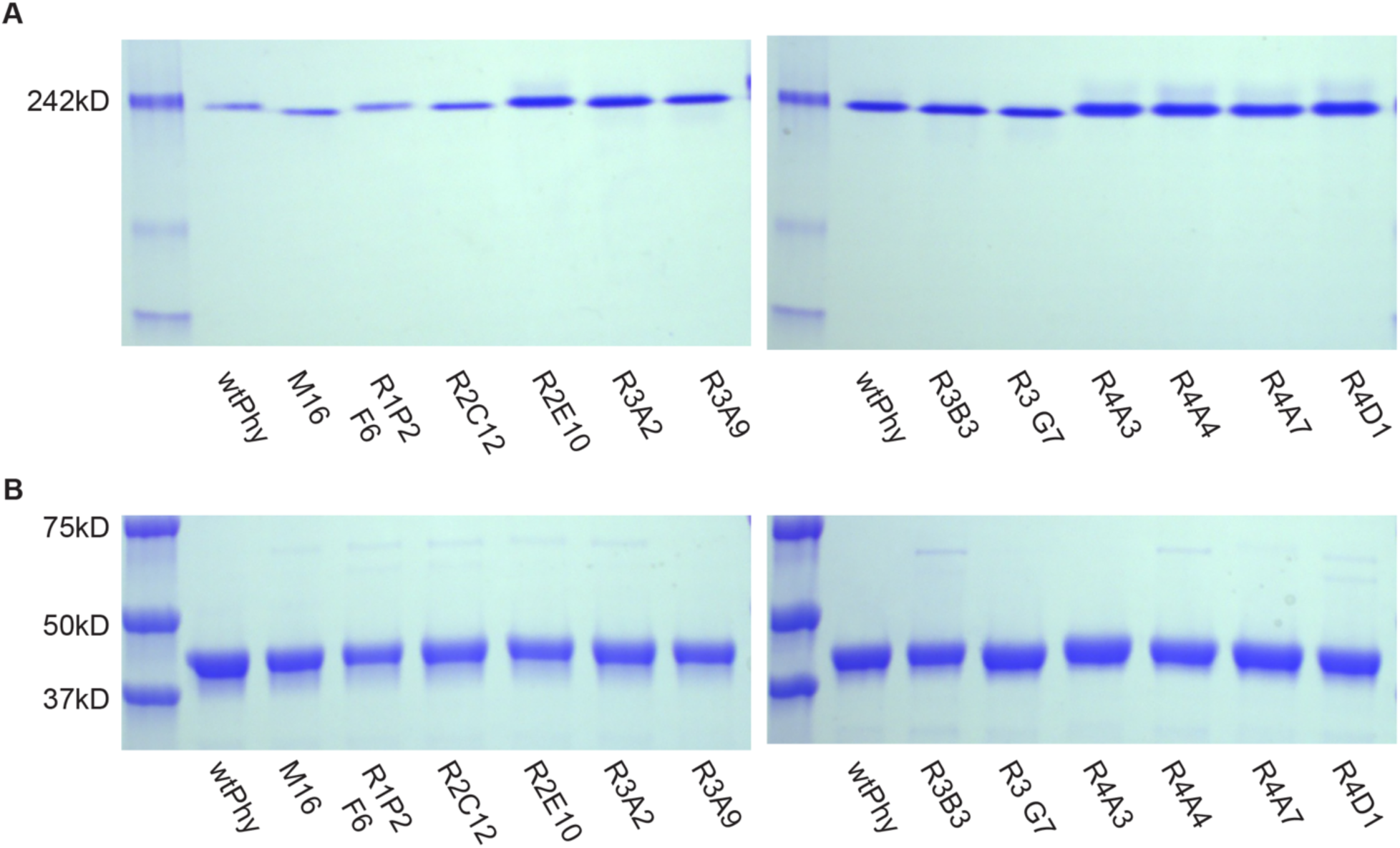
Native and SDS PAGE gel of purified *Ym*Phytase proteins. (**A**) For native PAGE, purified protein were mixed in native sample buffer and run in Tris/Glycine buffer at 7.5% TGX gel (BioRad #4568024) along with native protein ladder (Invitrogen # LC0725). As reported previously, the native gel confirmed that the phytase protein forms a tetramer. (**B**) For SDS-PAGE, purified proteins were mixed with 1x Laemmli buffer (+5% BME) and boiled at 95°C for five minutes. An equal amount of each sample was then loaded onto a 12% SDS-PAGE gel along with a protein ladder (BioRad # 1610374). After Coomassie staining, the gels were thoroughly washed before imaging.

**Extended Data Fig. 19.**
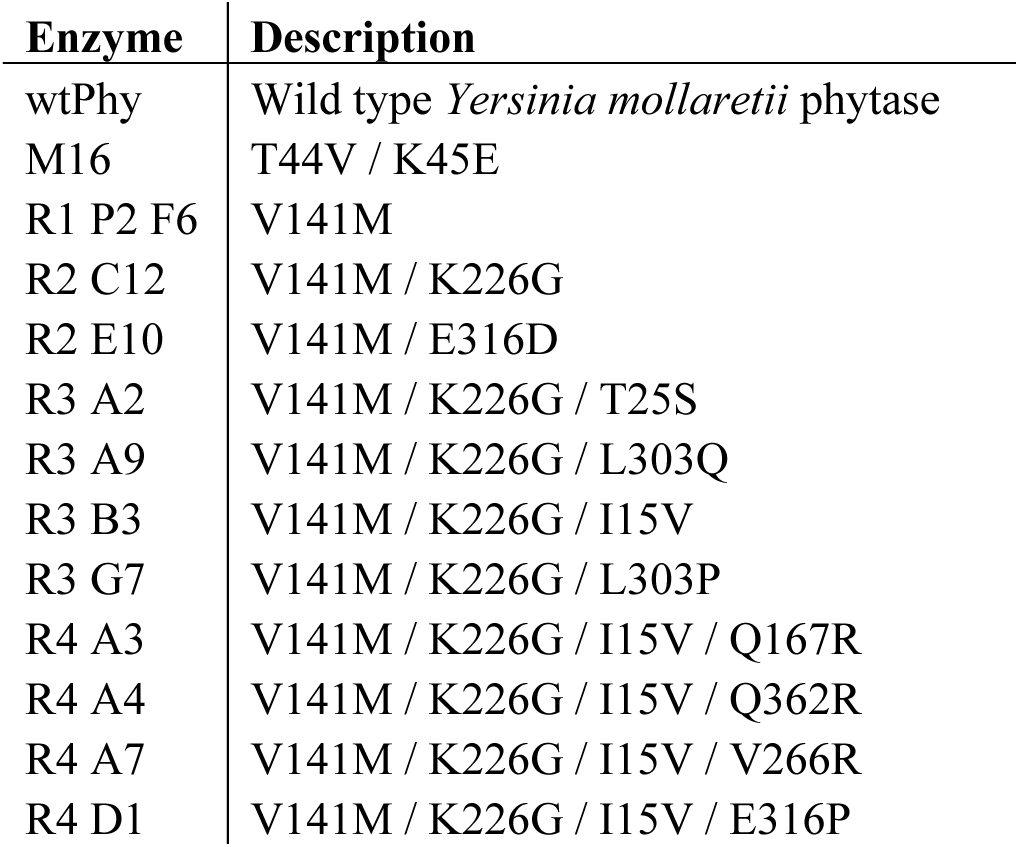
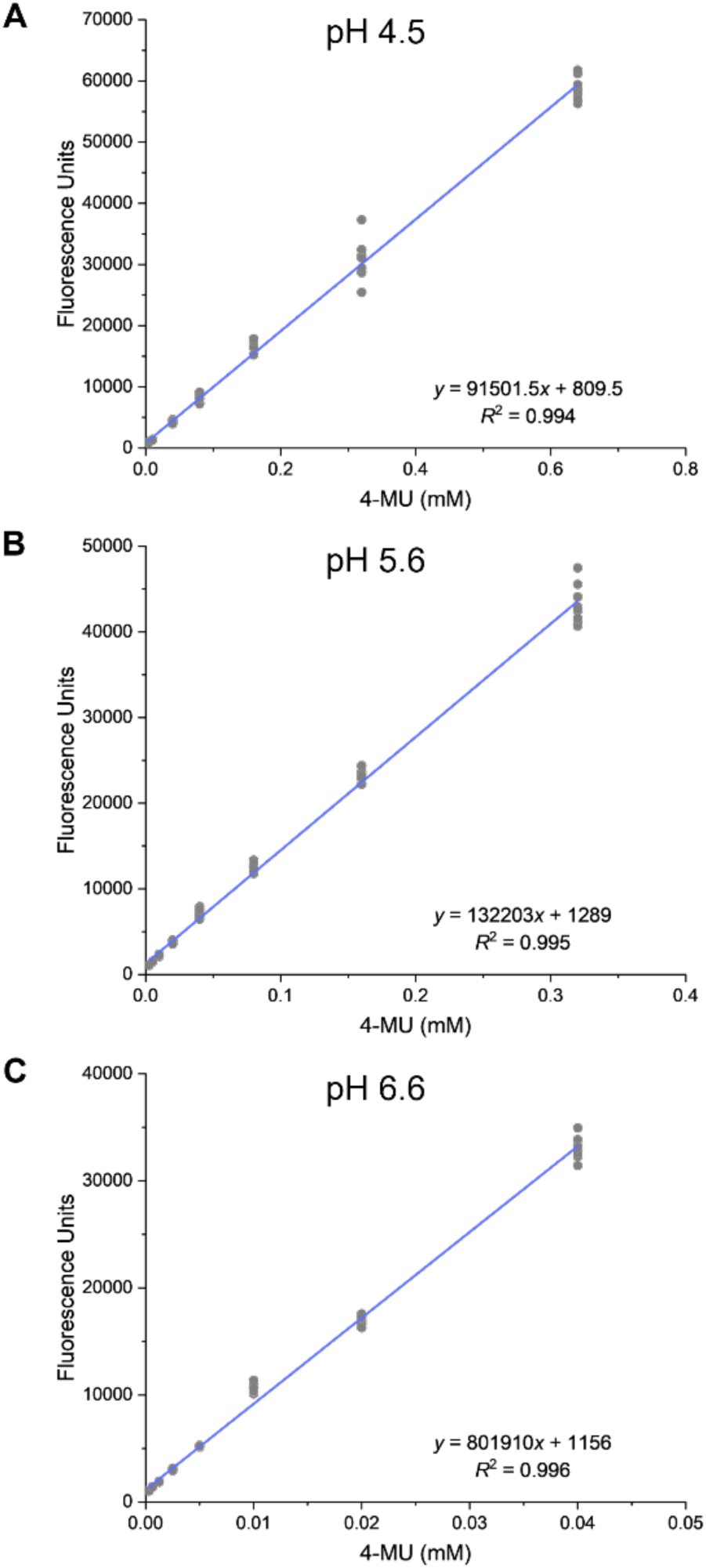
Standard curves for kinetic analysis of *Ym*Phytase at (A) pH 4.5, (B) pH 5.6, and (C) pH 6.6. Fluorescence values (*λ*_ex_ = 354 nm / *λ*_em_ = 465 nm) were correlated with 4-methylumbelliferone (4-MU) concentrations through making a standard curve. For pH 4.5, standards are comprised of eight different concentrations of 4-MU with eight independently prepared samples each, which are represented by grey dots. For pH 5.6 and 6.6, eight different concentrations of 4-MU were prepared in 12 independently prepared samples, which are represented by grey dots. 4-MU was dissolved into methanol and 50 µL of dissolved 4-MU solution was mixed with 50 µL of appropriate pH buffer. For pH 4.5, the buffer used was 0.25 M sodium acetate, 1 mM calcium chloride, 0.01% Tween-20 buffer. For pH 5.6 and 6.6, 0.2 M tris maleate buffer at the appropriate pH was used.

**Extended Data Fig. 20.**
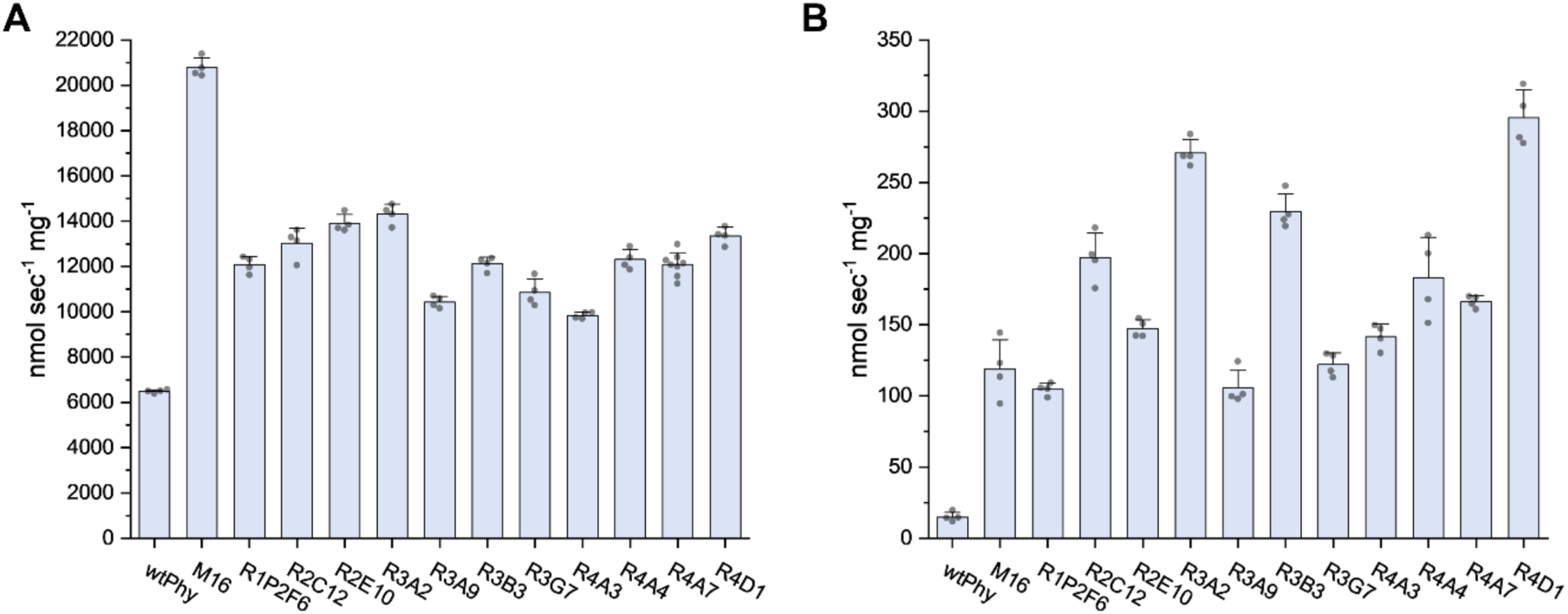
Specific enzyme activity of purified *Ym*Phytase variants at A) pH 4.5 and B) pH 5.6. Purified *Ym*Phytase variants were mixed with 1 mM AEBSF and incubated at 37 °C for 30 minutes prior to assays. Specific activity assays were performed with 0.9 mM 4-MUP. For pH 4.5, assays were performed in 0.25 M sodium acetate, 1 mM calcium chloride, and 0.01% Tween-20 buffer. For pH 5.6, assays were performed in 0.2 M tris maleate buffer. Bars indicate the average of four independent samples with standard deviations signified by error bars and individual data points as grey dots.

**Extended Data Fig. 21.**
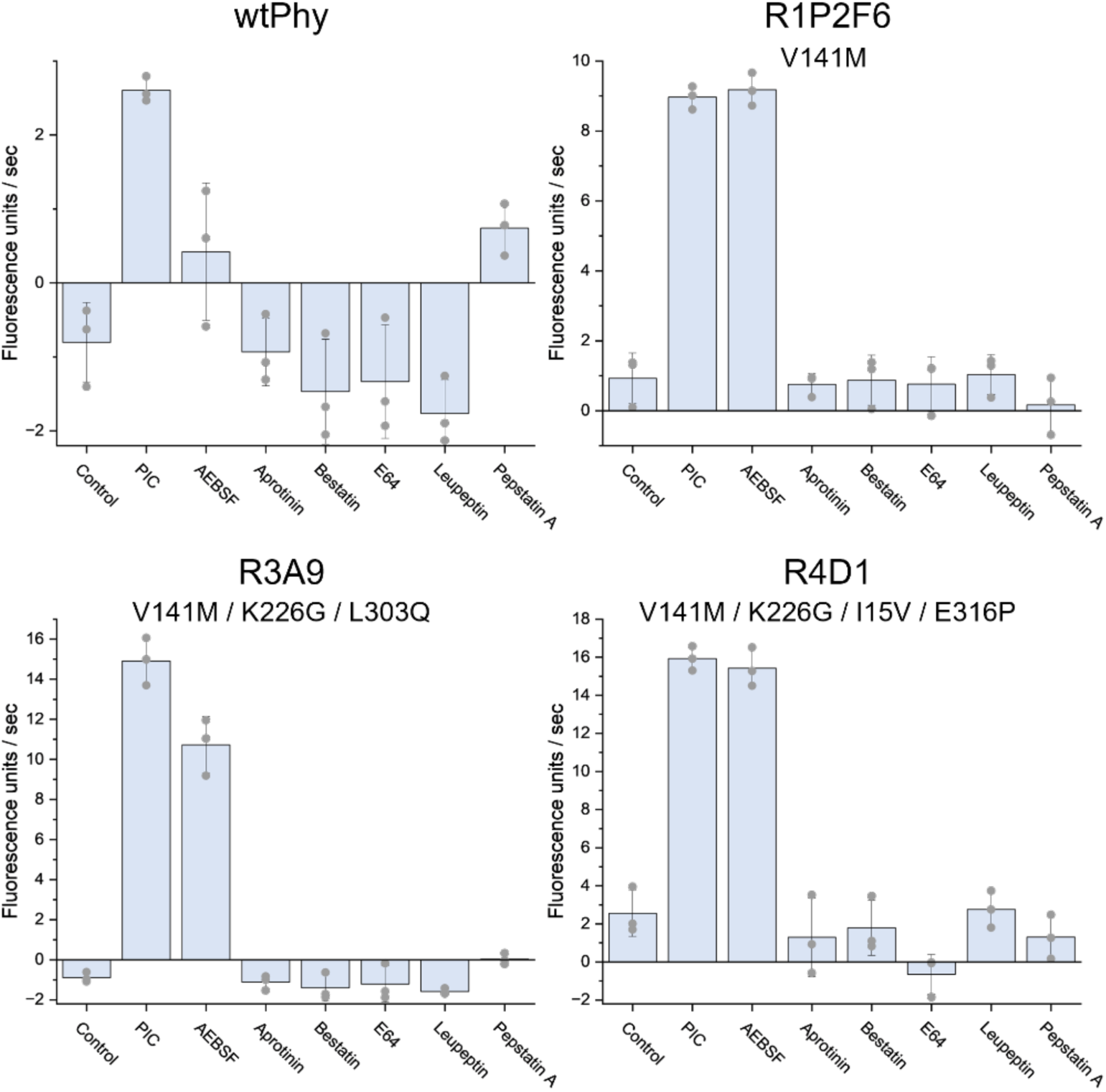
Effect of protease inhibitor additives to relative activity of purified phytase enzymes. Purified enzymes were mixed with Protease Inhibitor Cocktail (PIC, Sigma #P8849), 4-(2-aminoethyl)benzenesulfonyl fluoride hydrochloride (AEBSF), aprotinin, bestatin, protease inhibitor E64, leupeptin, or pepstatin A and incubated for 30 minutes at 37 °C, then used in a 4-MUP activity assay in 0.2 M tris maleate buffer at pH 6.6. Bars represent the average of three independent samples which are shown as grey dots.

**Extended Data Fig. 22.**
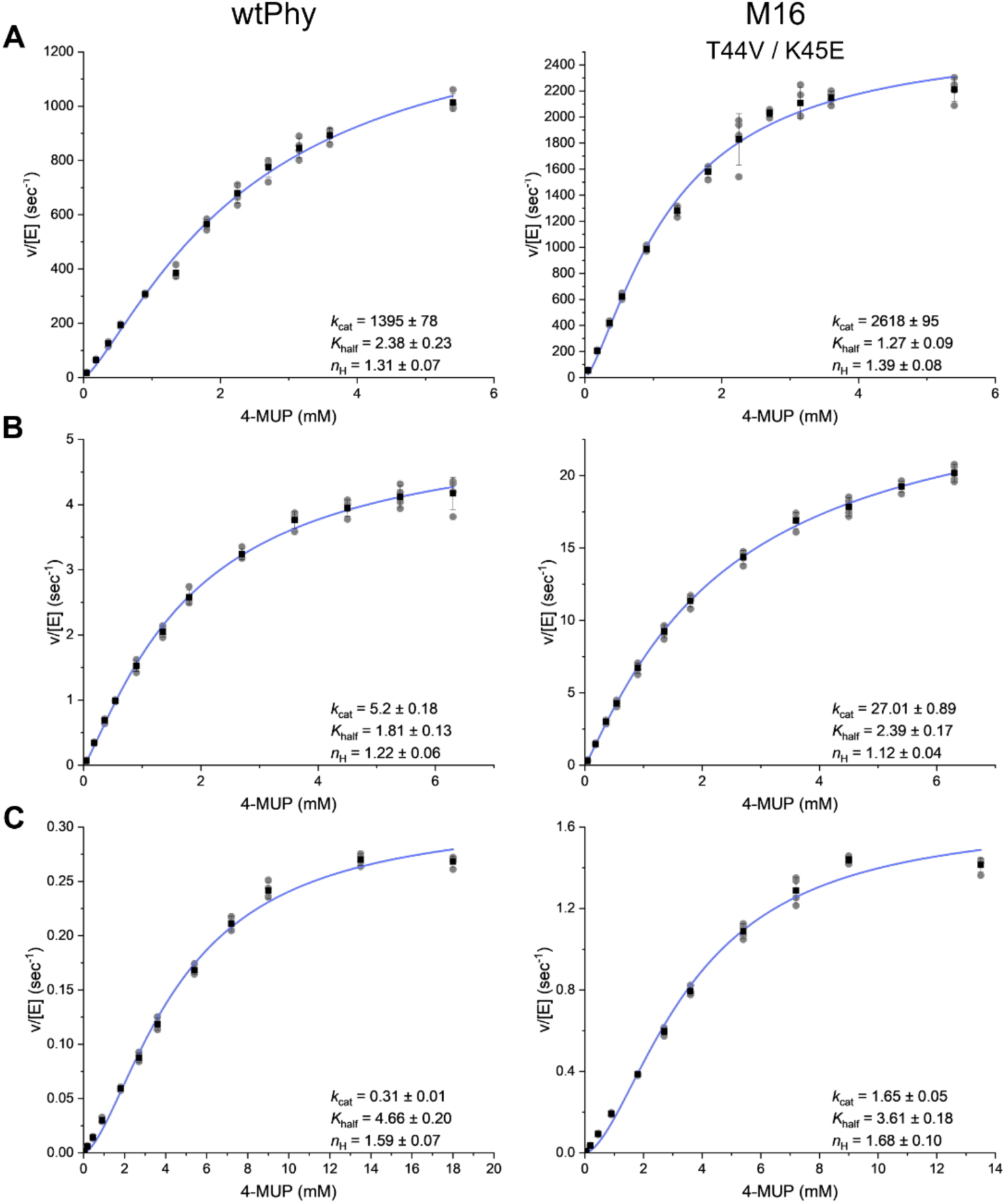
Kinetic analysis of wild type *Ym*Phytase (wtPhy, left) and positive control (M16, right) at (A) pH 4.5, (B) pH 5.6, and (C) pH 6.6. Purified phytase variants were mixed with 1 mM AEBSF and incubated at 37 °C for 30 minutes prior to kinetic assays. For pH 4.5, assays were performed in 0.25 M sodium acetate, 1 mM calcium chloride, and 0.01% Tween-20 buffer. For pH 5.6 and 6.6, assays were performed in 0.2 M tris maleate buffer. Black dots indicate the average of four independent samples with standard deviations signified by error bars and individual datapoints by grey dots.

**Extended Data Fig. 23.**
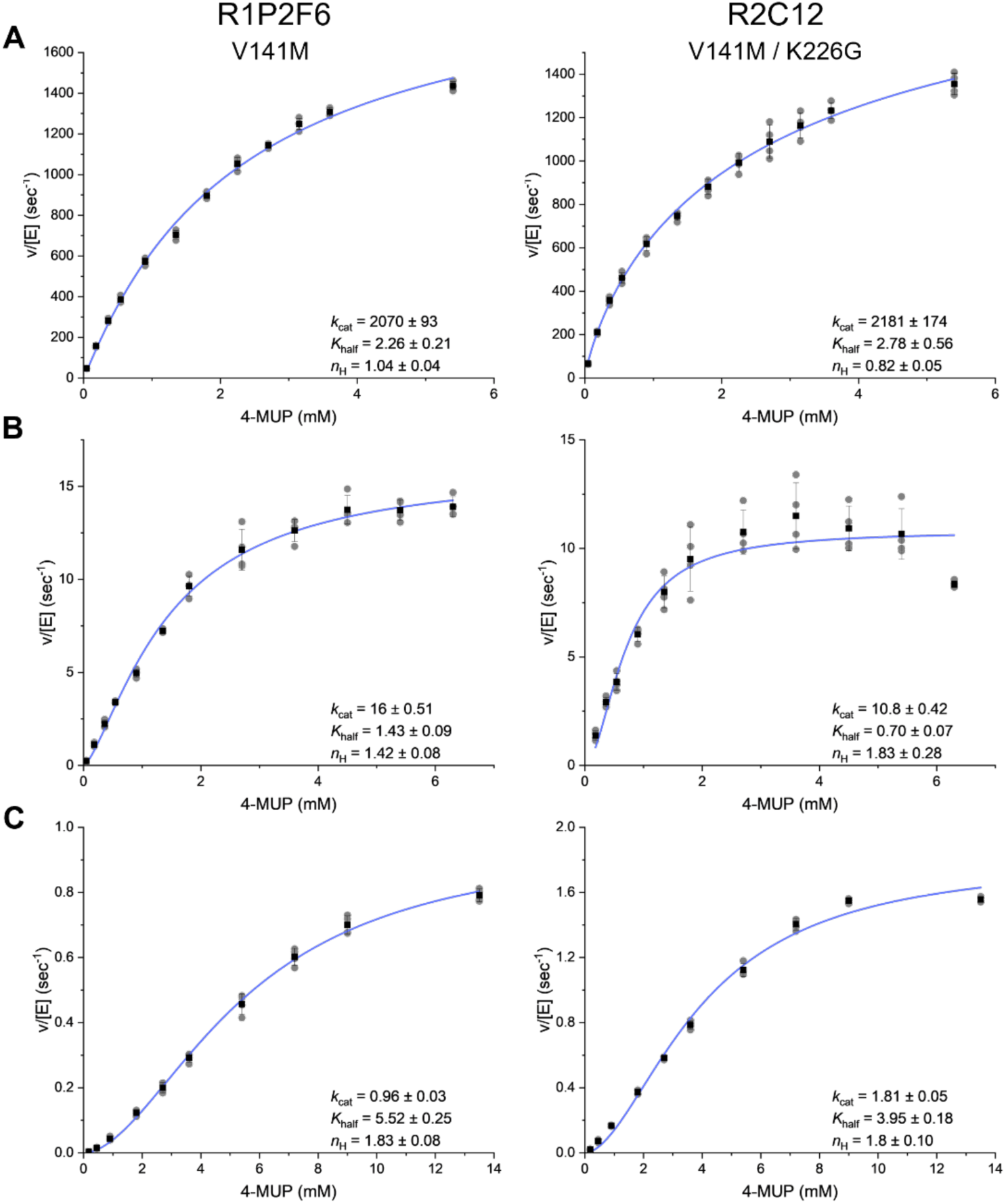
Kinetic analysis of *Ym*Phytase variants R1P2F6 (left) and R2C12 (right) at (A) pH 4.5, (B) pH 5.6, and (C) pH 6.6. Purified *Ym*Phytase variants were mixed with 1 mM AEBSF and incubated at 37 °C for 30 minutes prior to kinetic assays. For pH 4.5, assays were performed in 0.25 M sodium acetate, 1 mM calcium chloride, and 0.01% Tween-20 buffer. For pH 5.6 and 6.6, assays were performed in 0.2 M Tris maleate buffer. Black dots indicate the average of four independent samples with standard deviations signified by error bars and individual datapoints by grey dots.

**Extended Data Fig. 24.**
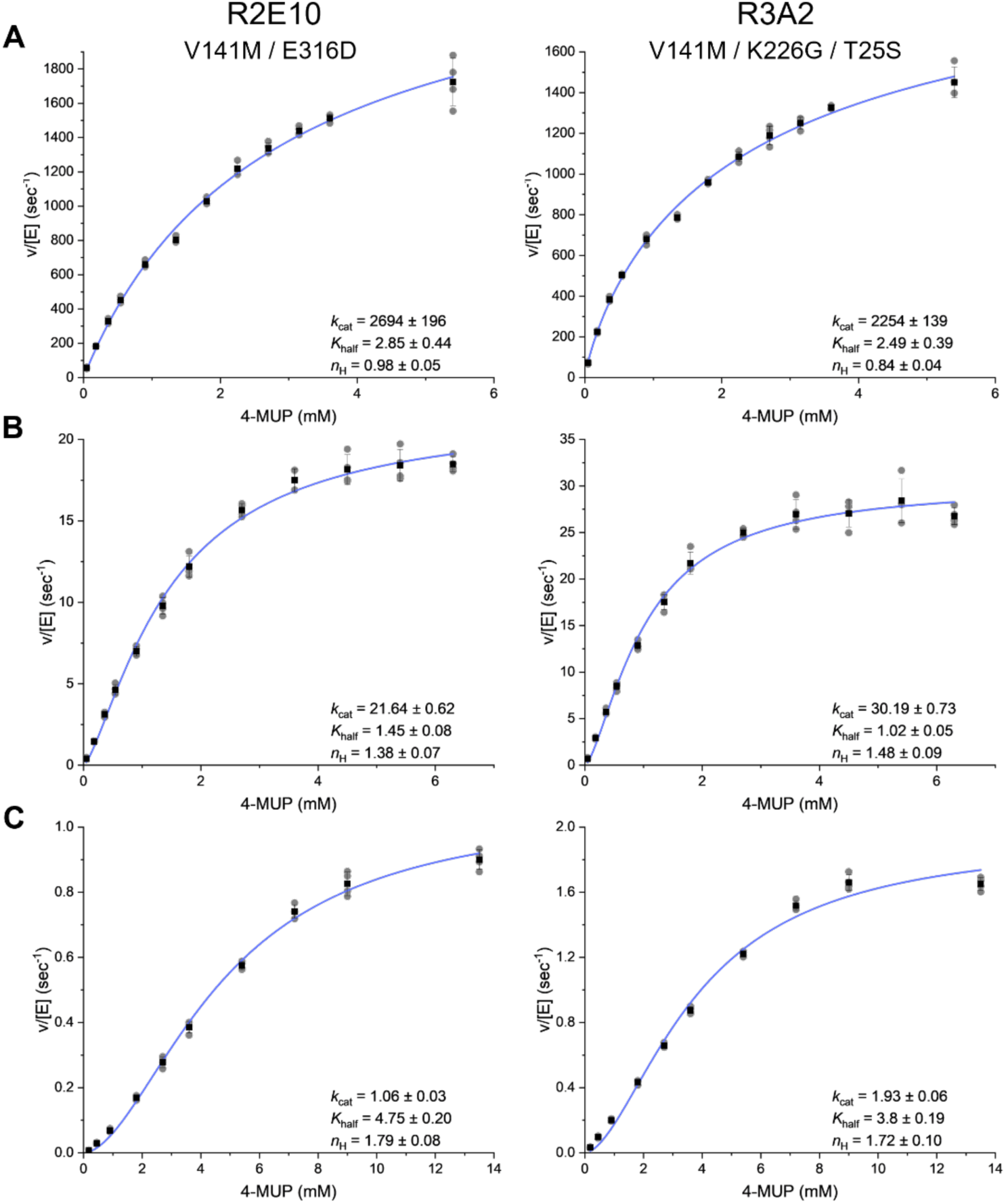
Kinetic analysis of *Ym*Phytase variants R2E10 (left) and R3A2 (right) at (A) pH 4.5, (B) pH 5.6, and (C) pH 6.6. Purified *Ym*Phytase variants were mixed with 1 mM AEBSF and incubated at 37 °C for 30 minutes prior to kinetic assays. For pH 4.5, assays were performed in 0.25 M sodium acetate, 1 mM calcium chloride, and 0.01% Tween-20 buffer. For pH 5.6 and 6.6, assays were performed in 0.2 M Tris maleate buffer. Black dots indicate the average of four independent samples with standard deviations signified by error bars and individual datapoints by grey dots.

**Extended Data Fig. 25.**
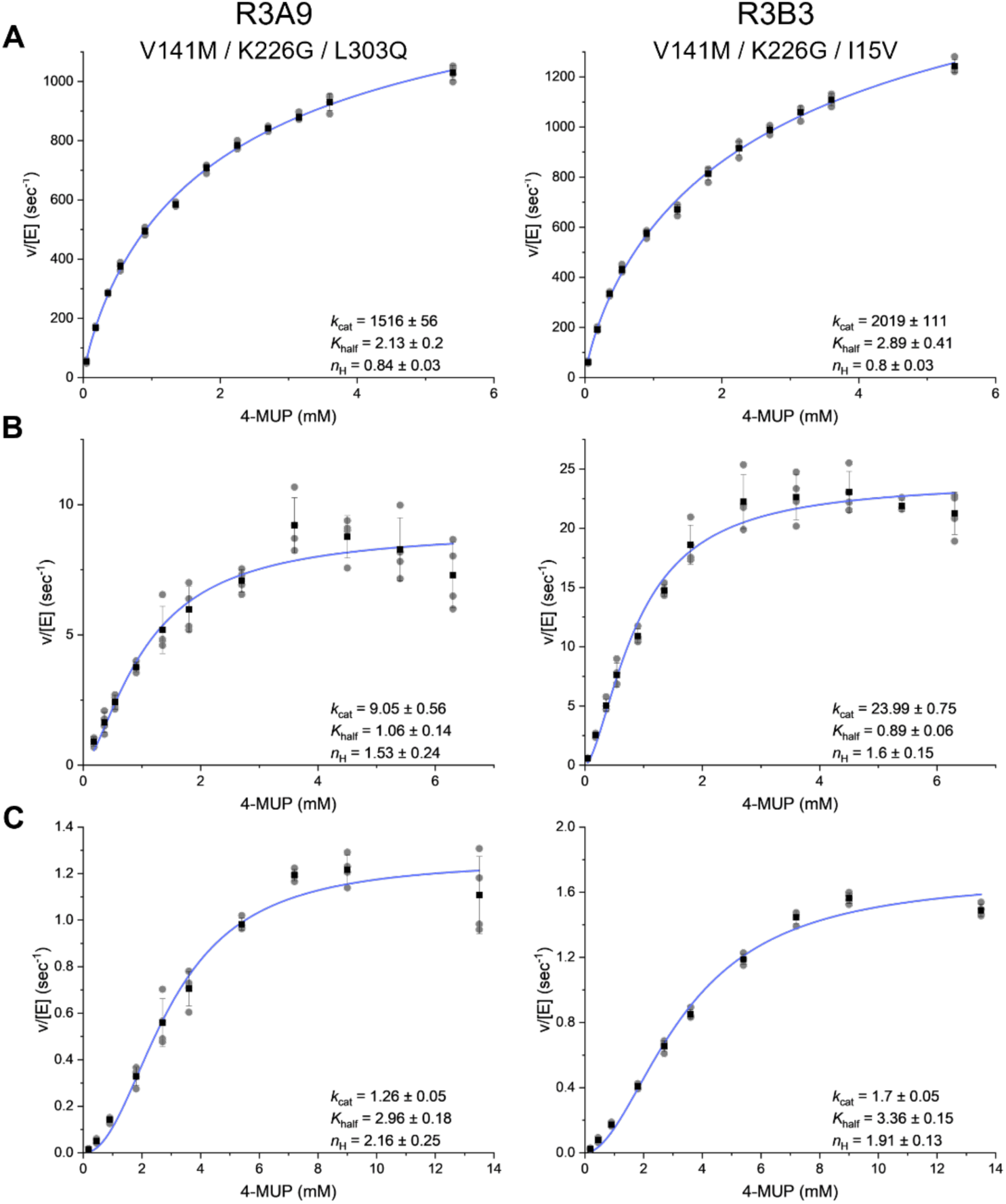
Kinetic analysis of *Ym*Phytase variants R3A9 (left) and R3B3 (right) at (A) pH 4.5, (B) pH 5.6, and (C) pH 6.6. Purified *Ym*Phytase variants were mixed with 1 mM AEBSF and incubated at 37 °C for 30 minutes prior to kinetic assays. For pH 4.5, assays were performed in 0.25 M sodium acetate, 1 mM calcium chloride, and 0.01% Tween-20 buffer. For pH 5.6 and 6.6, assays were performed in 0.2 M Tris maleate buffer. Black dots indicate the average of four independent samples with standard deviations signified by error bars and individual datapoints by grey dots.

**Extended Data Fig. 26.**
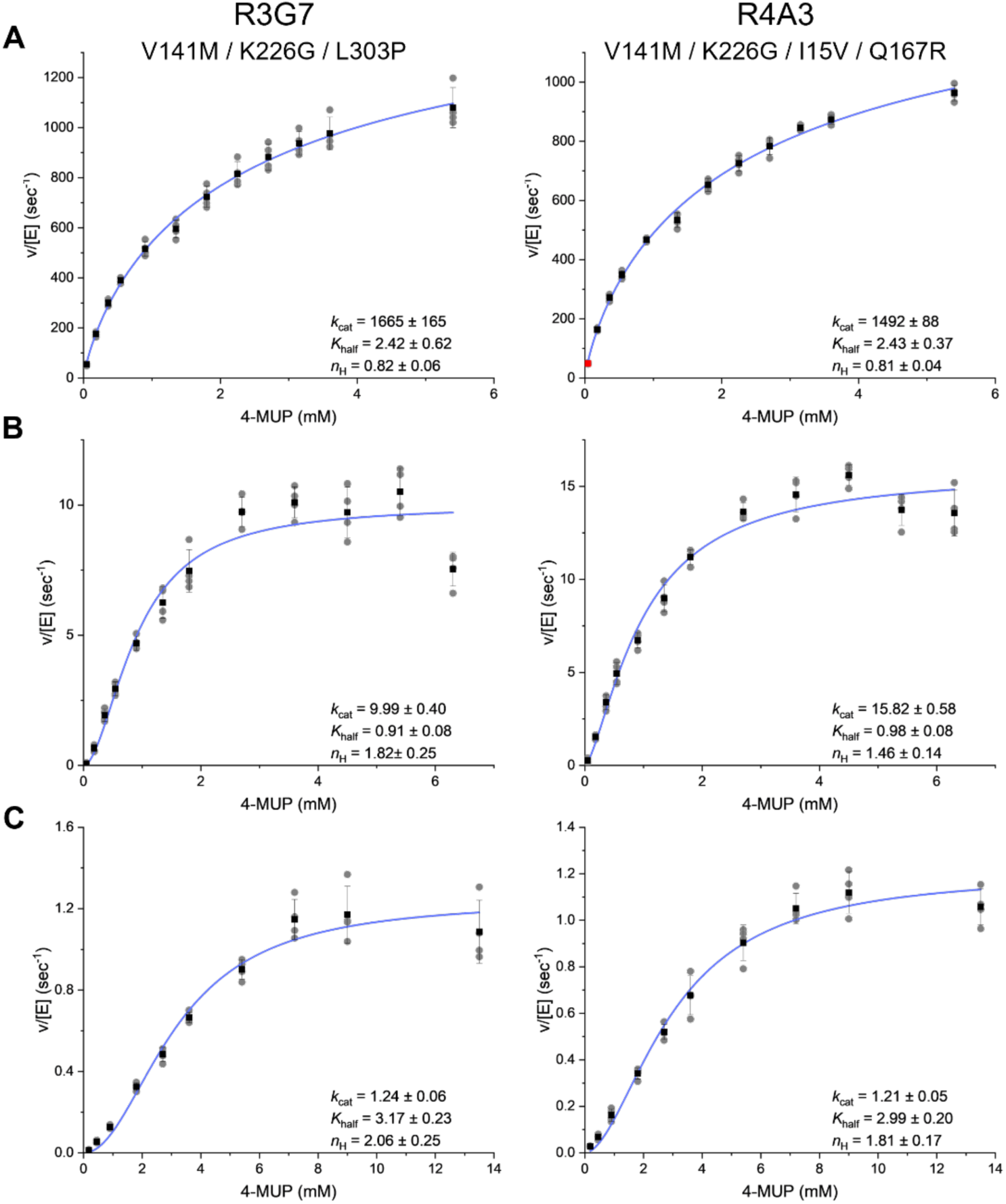
Kinetic characterization of *Ym*Phytase variants R3G7 (left) and R4A3 (right) at (A) pH 4.5, (B) pH 5.6, and (C) pH 6.6. Purified *Ym*Phytase variants were mixed with 1 mM AEBSF and incubated at 37 °C for 30 minutes prior to kinetic assays. For pH 4.5, assays were performed in 0.25 M sodium acetate, 1 mM calcium chloride, and 0.01% Tween-20 buffer. For pH 5.6 and 6.6, assays were performed in 0.2 M Tris maleate buffer. Black dots indicate the average of four independent samples with standard deviations signified by error bars and individual datapoints by grey dots.

**Extended Data Fig. 27.**
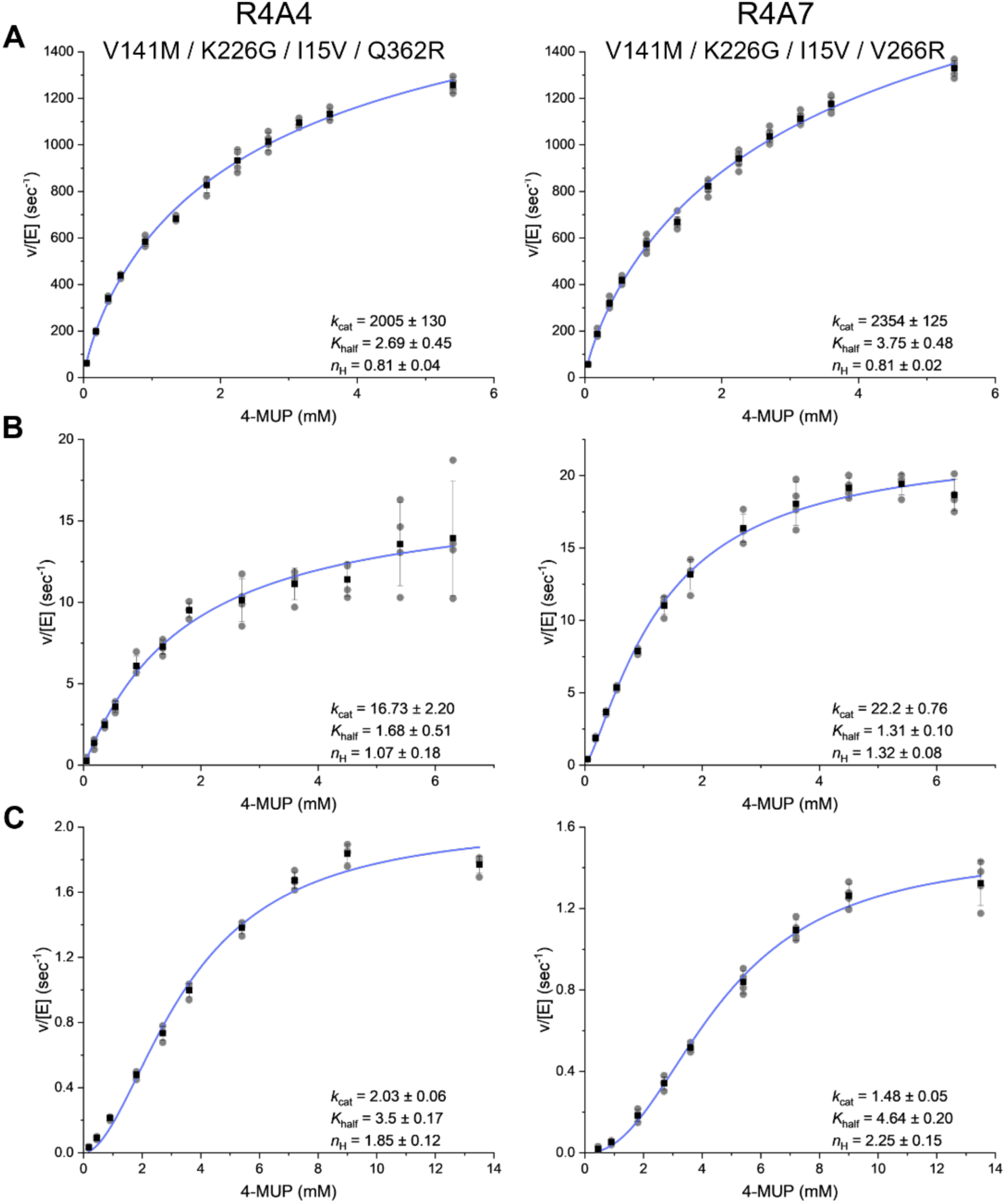
Kinetic characterization of *Ym*Phytase variants R4A4 (left) and R4A7 (right) at (A) pH 4.5, (B) pH 5.6, and (C) pH 6.6. Purified *Ym*Phytase variants were mixed with 1 mM AEBSF and incubated at 37 °C for 30 minutes prior to kinetic assays. For pH 4.5, assays were performed in 0.25 M sodium acetate, 1 mM calcium chloride, and 0.01% Tween-20 buffer. For pH 5.6 and 6.6, assays were performed in 0.2 M Tris maleate buffer. Black dots indicate the average of four independent samples with standard deviations signified by error bars and individual datapoints by grey dots.

**Extended Data Fig. 28.**
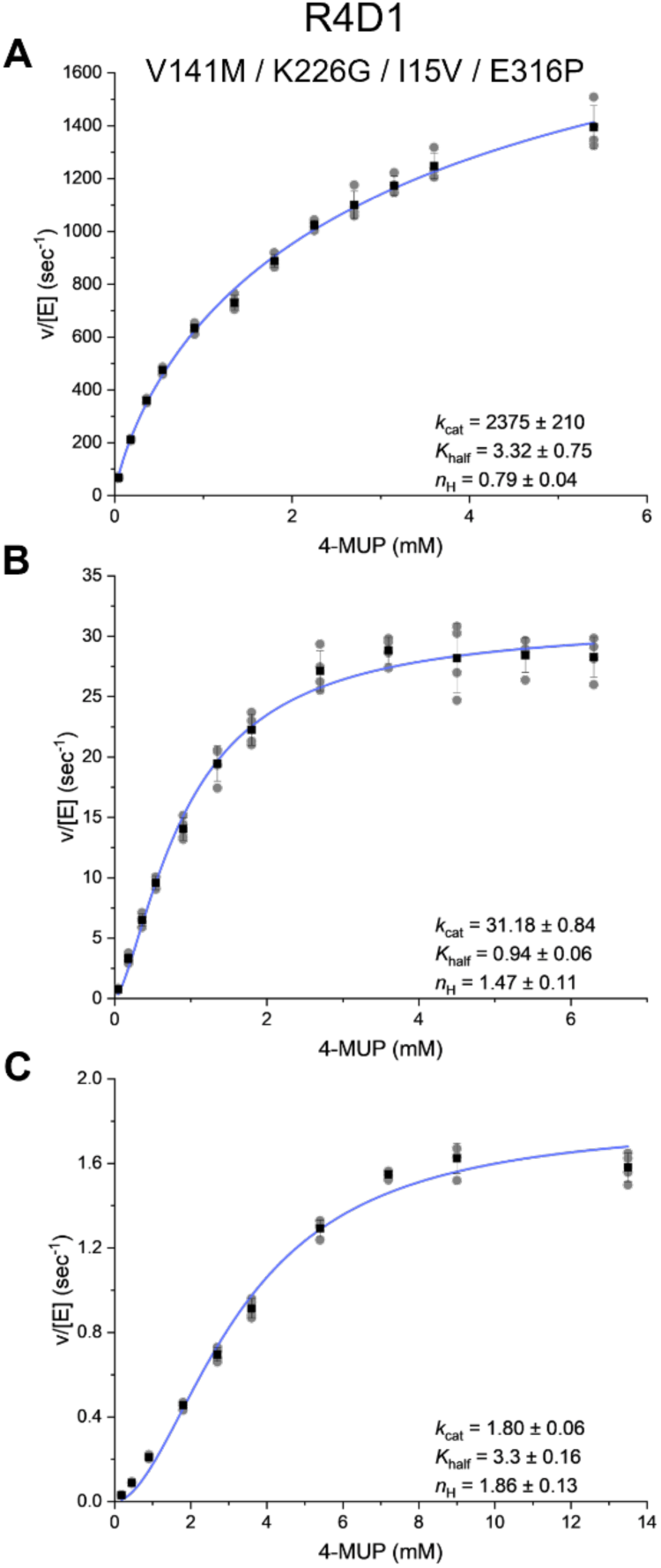
Kinetic characterization of *Ym*Phytase variant R4D1 at (A) pH 4.5, (B) pH 5.6, and (C) pH 6.6. Purified phytase variants were mixed with 1 mM AEBSF and incubated at 37 °C for 30 minutes prior to kinetic assays. For pH 4.5, assays were performed in 0.25 M sodium acetate, 1 mM calcium chloride, and 0.01% Tween-20 buffer. For pH 5.6 and 6.6, assays were performed in 0.2 M Tris maleate buffer. Black dots indicate the average of four independent samples with standard deviations signified by error bars and individual datapoints by grey dots.

**Extended Data Fig. 29.**
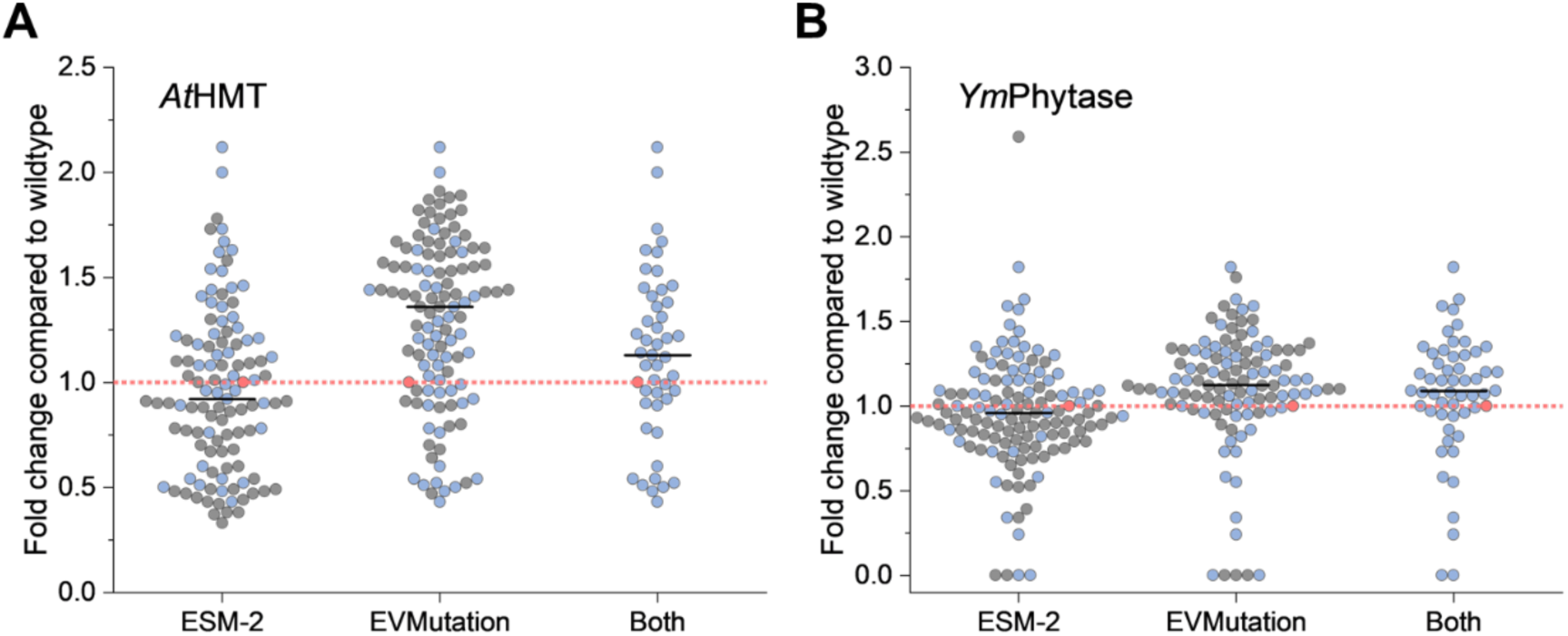
Comparison of models used for prediction of initial variants of (**A**) *At*HMT and **(B)** *Ym*Phytase. Circles indicate variant performance relative to the wild-type enzyme (pink circle and dotted line). Mutants ranked within the top 200 predictions of both ESM-2 and EVmutation are pictured as blue circles. Mutants predicted solely by the indicated model or ranked lower than 200 by the opposing model are shown as grey circles. The black line indicates the median of all samples within the column. For *At*HMT, 26.1% (46 of 176 total predictions) of variants were predicted by both ESM-2 and EVmutation. For *Ym*Phytase, 29.4% (53 of 180 total predictions) of variants were predicted by both models.

**Extended Data Table 1.** Michaelis-Menten kinetics of wild type *At*HMT and its variants using ethyl iodide as a substrate.

| HMT variant | $K_m$ | $k_{cat}$ | Mutations |
| --- | --- | --- | --- |
| wtHMT | $37.75 \pm 3.74$ | $0.47 \pm 0.02$ | Wild type HMT from *Arabidopsis thaliana* |
| V140T | $25.95 \pm 2.37$ | $1.53 \pm 0.06$ | V140T |
| R2 A1 | $19.75 \pm 4.66$ | $1.56 \pm 0.15$ | V140T / S99T |
| R2 C3 | $27.50 \pm 4.28$ | $1.93 \pm 0.13$ | V140T / D9A |
| R3 A1 | $13.81 \pm 2.45$ | $1.50 \pm 0.09$ | V140T / L101I / D9A |
| R3 A2 | $13.93 \pm 2.71$ | $1.68 \pm 0.13$ | V140T / L101I / V195H |
| R3 A4 | $11.39 \pm 2.52$ | $1.36 \pm 0.1$ | V140T / L101I / D9Q |
| R3 C7 | $17.74 \pm 2.2$ | $1.71 \pm 0.08$ | V140T / L101I / T107L |
| R4 A1 | $19.89 \pm 5.31$ | $3.63 \pm 0.38$ | V140T / L101I / T213S / V178E |
| R4 B2 | $20.81 \pm 4.69$ | $3.01 \pm 0.27$ | V140T / L101I / S63N / E32N |
| R4 B3 | $26.91 \pm 4.55$ | $4.41 \pm 0.32$ | V140T / L101I / V178E / E32N |
| R4 C2 | $19.69 \pm 4.79$ | $4.84 \pm 0.46$ | V140T / L101I / E206D / S63N |

\*The notation R4 C2 signifies well C2 from the fourth round of evolution.

**Extended Data Table 2.**
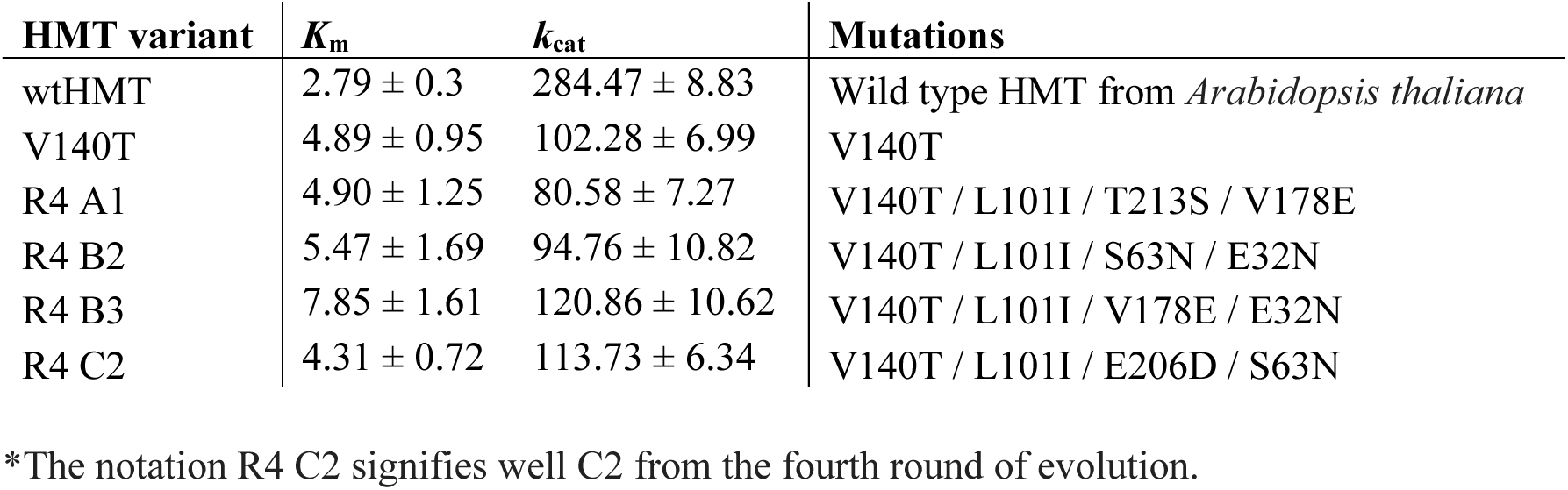
Michaelis-Menten kinetics of wild type *At*HMT and its variants using methyl iodide as a substrate.

| **HMT variant** | **$K_m$** | **$k_{cat}$** | **Mutations** |
| --- | --- | --- | --- |
| wtHMT | $2.79 \pm 0.3$ | $284.47 \pm 8.83$ | Wild type HMT from *Arabidopsis thaliana* |
| V140T | $4.89 \pm 0.95$ | $102.28 \pm 6.99$ | V140T |
| R4 A1 | $4.90 \pm 1.25$ | $80.58 \pm 7.27$ | V140T / L101I / T213S / V178E |
| R4 B2 | $5.47 \pm 1.69$ | $94.76 \pm 10.82$ | V140T / L101I / S63N / E32N |
| R4 B3 | $7.85 \pm 1.61$ | $120.86 \pm 10.62$ | V140T / L101I / V178E / E32N |
| R4 C2 | $4.31 \pm 0.72$ | $113.73 \pm 6.34$ | V140T / L101I / E206D / S63N |

\*The notation R4 C2 signifies well C2 from the fourth round of evolution.

**Extended Data Table 3.** Specific activities of wild type *At*HMT and its variants using ethyl iodide a substrate.

| HMT variant | Specific activity<br>(nmol min<sup>-1</sup> mg<sup>-1</sup>) | Mutations |
| --- | --- | --- |
| wtHMT | 4.72 ± 0.42 | Wild type HMT from *Arabidopsis thaliana* |
| V140T | 19.78 ± 0.72 | V140T |
| R2A1 | 19.94 ± 3.56 | V140T / S99T |
| R2C3 | 21.27 ± 2.31 | V140T / D9A |
| R3A1 | 26.87 ± 2.63 | V140T / L101I / D9A |
| R3A2 | 36.12 ± 0.72 | V140T / L101I / V195H |
| R3A4 | 34.97 ± 5.32 | V140T / L101I / D9Q |
| R3C7 | 30.15 ± 0.50 | V140T / L101I / T107L |
| R4A1 | 62.02 ± 6.06 | V140T / L101I / T213S / V178E |
| R4B2 | 37.63 ± 7.06 | V140T / L101I / S63N / E32N |
| R4B3 | 64.63 ± 6.52 | V140T / L101I / V178E / E32N |
| R4C2 | 79.57 ± 2.57 | V140T / L101I / E206D / S63N |

**Extended Data Table 4.** Substrate preference of wild type *At*HMT and its variants for ethyl iodide (EtI) and methyl iodide (MeI).

| HMT<br>variant | $k_{\text{cat}}/K_{\text{m}}$ | | $k_{\text{cat}}/K_{\text{m}}$<br>ratio | Relative<br>substrate<br>preference<br>(EtI/MeI) | Mutations |
| --- | --- | --- | --- | --- | --- |
|  | Ethyl<br>iodide | Methyl<br>iodide |  |  |  |
| wtHMT | 0.01 | 102.34 | 1.22E-04 | 1 | Wild type HMT from *Arabidopsis thaliana* |
| V140T | 0.06 | 20.92 | 2.80E-03 | 23.01 | V140T |
| R4 A1 | 0.18 | 16.44 | 1.11E-02 | 91.2 | V140T / L101I / T213S / V178E |
| R4 B2 | 0.14 | 17.32 | 8.35E-03 | 68.64 | V140T / L101I / S63N / E32N |
| R4 B3 | 0.16 | 15.4 | 1.06E-02 | 87.47 | V140T / L101I / V178E / E32N |
| R4 C2 | 0.25 | 26.39 | 9.32E-03 | 76.55 | V140T / L101I / E206D / S63N |

\*The relative substrate preference is calculated as fold change, with respect to the wtHMT, in the ratio of $k_{\text{cat}}/K_{\text{m}}$ for ethyl iodide and methyl iodide.

**Extended Data Table 5.**
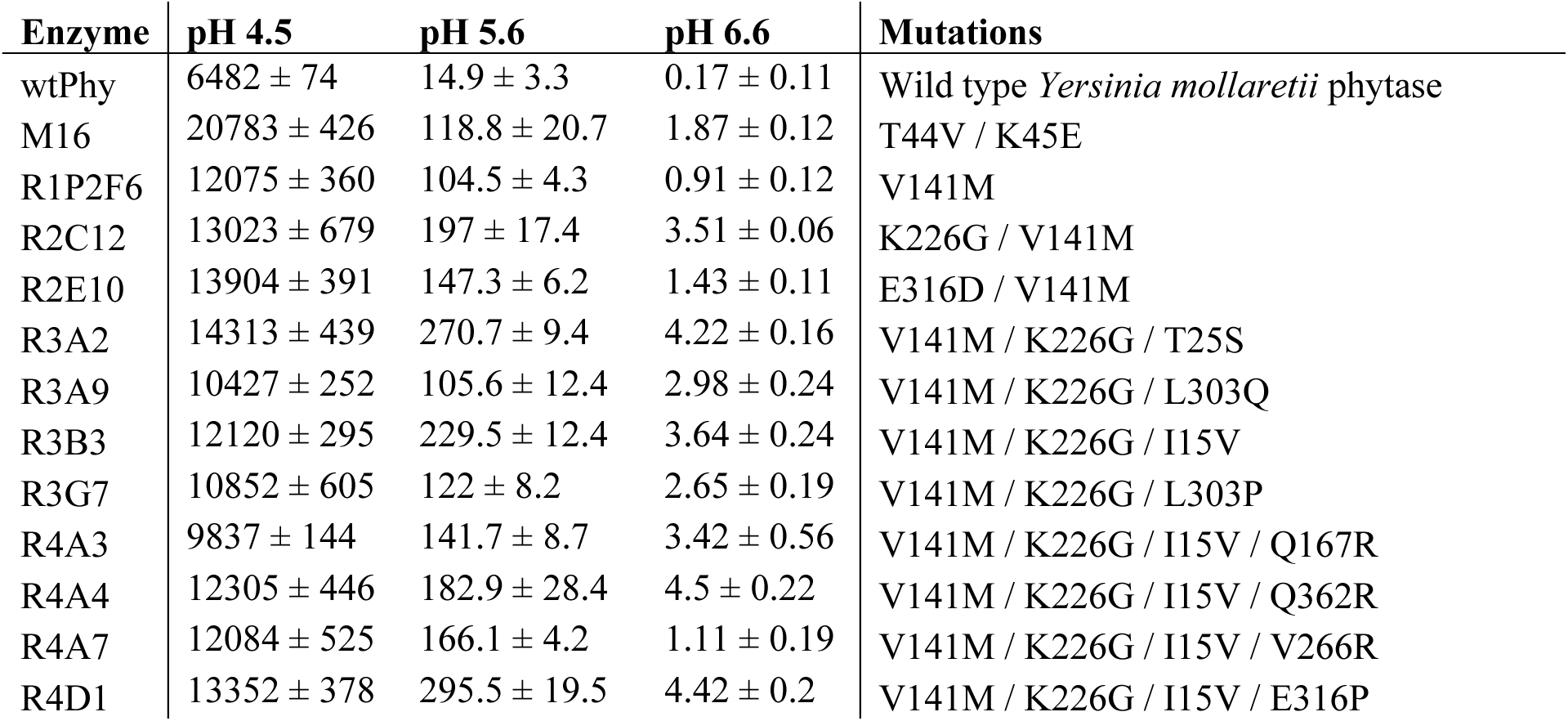
Specific activities of wild type *Ym*Phytase and its variants. The specific activity was determined at pH 4.5, 5.6, and 6.6 with a substrate concentration of 0.9 mM 4-MUP. Displayed values are the average of at least four independent samples with standard deviations. Specific activities are expressed as nmol 4-MU / second / mg phytase.

| Enzyme | pH 4.5 | pH 5.6 | pH 6.6 | Mutations |
| --- | --- | --- | --- | --- |
| wtPhy | 6482 ± 74 | 14.9 ± 3.3 | 0.17 ± 0.11 | Wild type *Yersinia mollaretii* phytase |
| M16 | 20783 ± 426 | 118.8 ± 20.7 | 1.87 ± 0.12 | T44V / K45E |
| R1P2F6 | 12075 ± 360 | 104.5 ± 4.3 | 0.91 ± 0.12 | V141M |
| R2C12 | 13023 ± 679 | 197 ± 17.4 | 3.51 ± 0.06 | K226G / V141M |
| R2E10 | 13904 ± 391 | 147.3 ± 6.2 | 1.43 ± 0.11 | E316D / V141M |
| R3A2 | 14313 ± 439 | 270.7 ± 9.4 | 4.22 ± 0.16 | V141M / K226G / T25S |
| R3A9 | 10427 ± 252 | 105.6 ± 12.4 | 2.98 ± 0.24 | V141M / K226G / L303Q |
| R3B3 | 12120 ± 295 | 229.5 ± 12.4 | 3.64 ± 0.24 | V141M / K226G / I15V |
| R3G7 | 10852 ± 605 | 122 ± 8.2 | 2.65 ± 0.19 | V141M / K226G / L303P |
| R4A3 | 9837 ± 144 | 141.7 ± 8.7 | 3.42 ± 0.56 | V141M / K226G / I15V / Q167R |
| R4A4 | 12305 ± 446 | 182.9 ± 28.4 | 4.5 ± 0.22 | V141M / K226G / I15V / Q362R |
| R4A7 | 12084 ± 525 | 166.1 ± 4.2 | 1.11 ± 0.19 | V141M / K226G / I15V / V266R |
| R4D1 | 13352 ± 378 | 295.5 ± 19.5 | 4.42 ± 0.2 | V141M / K226G / I15V / E316P |

**Extended Data Table 6.** Enzyme kinetics of wild type *Ym*Phytase and its variants using the 4-methylumbelliferyl phosphate (4-MUP) assay.

| Enzyme | pH 4.5 |  |  | pH 5.6 |  |  | pH 6.6 |  |  |
| --- | --- | --- | --- | --- | --- | --- | --- | --- | --- |
| | $k_{cat}$ | $K_{half}$ | $n_H$ | $k_{cat}$ | $K_{half}$ | $n_H$ | $k_{cat}$ | $K_{half}$ | $n_H$ |
| wtPhy | 1396 ± 78 | 2.38 ± 0.23 | 1.31 ± 0.07 | 5.2 ± 0.18 | 1.81 ± 0.13 | 1.22 ± 0.06 | 0.31 ± 0.01 | 4.66 ± 0.2 | 1.59 ± 0.07 |
| M16 | 2653 ± 74 | 1.27 ± 0.07 | 1.41 ± 0.06 | 27.01 ± 0.89 | 2.39 ± 0.17 | 1.12 ± 0.04 | 1.65 ± 0.05 | 3.61 ± 0.18 | 1.68 ± 0.1 |
| R1P2F6 | 2071 ± 93 | 2.26 ± 0.21 | 1.04 ± 0.04 | 16 ± 0.51 | 1.43 ± 0.09 | 1.42 ± 0.08 | 0.96 ± 0.03 | 5.52 ± 0.25 | 1.83 ± 0.08 |
| R2C12 | 2182 ± 174 | 2.78 ± 0.56 | 0.82 ± 0.05 | 10.8 ± 0.42 | 0.7 ± 0.07 | 1.83 ± 0.28 | 1.81 ± 0.05 | 3.95 ± 0.18 | 1.8 ± 0.1 |
| R2E10 | 2089 ± 107 | 1.74 ± 0.18 | 1.13 ± 0.06 | 21.64 ± 0.62 | 1.45 ± 0.08 | 1.38 ± 0.07 | 1.06 ± 0.03 | 4.75 ± 0.2 | 1.79 ± 0.08 |
| R3A2 | 2176 ± 118 | 2.29 ± 0.31 | 0.85 ± 0.04 | 30.19 ± 0.73 | 1.02 ± 0.05 | 1.48 ± 0.09 | 1.93 ± 0.06 | 3.8 ± 0.19 | 1.72 ± 0.1 |
| R3A9 | 1517 ± 56 | 2.13 ± 0.2 | 0.84 ± 0.03 | 9.05 ± 0.56 | 1.06 ± 0.14 | 1.53 ± 0.24 | 1.26 ± 0.05 | 2.96 ± 0.18 | 2.16 ± 0.25 |
| R3B3 | 2020 ± 111 | 2.89 ± 0.41 | 0.8 ± 0.03 | 23.99 ± 0.75 | 0.89 ± 0.06 | 1.6 ± 0.15 | 1.7 ± 0.05 | 3.36 ± 0.15 | 1.91 ± 0.13 |
| R3G7 | 1666 ± 165 | 2.42 ± 0.62 | 0.82 ± 0.06 | 9.99 ± 0.4 | 0.91 ± 0.08 | 1.82 ± 0.25 | 1.24 ± 0.06 | 3.17 ± 0.23 | 2.06 ± 0.25 |
| R4A3 | 1493 ± 88 | 2.43 ± 0.37 | 0.81 ± 0.04 | 15.82 ± 0.58 | 0.98 ± 0.08 | 1.46 ± 0.14 | 1.21 ± 0.05 | 2.99 ± 0.2 | 1.81 ± 0.17 |
| R4A4 | 2006 ± 130 | 2.69 ± 0.45 | 0.81 ± 0.04 | 16.73 ± 2.2 | 1.68 ± 0.51 | 1.07 ± 0.18 | 2.03 ± 0.06 | 3.5 ± 0.17 | 1.85 ± 0.12 |
| R4A7 | 2355 ± 125 | 3.75 ± 0.48 | 0.81 ± 0.02 | 22.2 ± 0.76 | 1.31 ± 0.1 | 1.32 ± 0.08 | 1.48 ± 0.05 | 4.64 ± 0.2 | 2.25 ± 0.15 |
| R4D1 | 2376 ± 210 | 3.32 ± 0.75 | 0.79 ± 0.04 | 31.18 ± 0.84 | 0.94 ± 0.06 | 1.47 ± 0.11 | 1.8 ± 0.06 | 3.3 ± 0.16 | 1.86 ± 0.13 |

\*An enzyme label of R4D1 signifies the variant in well D1 from the fourth round of autonomous protein engineering. Mutations present in the variants listed above can be found in Table S5.

